# A masculinizing supergene underlies an exaggerated male reproductive morph in a spider

**DOI:** 10.1101/2021.02.09.430505

**Authors:** Frederik Hendrickx, Zoë De Corte, Gontran Sonet, Steven M. Van Belleghem, Stephan Köstlbacher, Carl Vangestel

## Abstract

In many species, individuals can develop into strikingly different morphs, which are determined by a simple Mendelian locus. How selection shapes loci that control complex phenotypic differences remains poorly understood. In the spider *Oedothorax gibbosus*, males either develop into a ‘hunched’ morph with conspicuous head structures or as a fast developing ‘flat’ morph with a female-like appearance. We show that the hunched-differs from the flat-determining allele by a hunch-specific genomic fragment of approximately 3 megabases. This fragment comprises dozens of genes that duplicated from genes found at different chromosomes. All functional duplicates, including *doublesex* - a key sexual differentiation regulatory gene, show male-specific expression, which illustrates their combined role as a masculinizing supergene. Our findings demonstrate how extensive indel polymorphisms and duplications of regulatory genes may contribute to the evolution of co-adapted gene clusters, sex-limited reproductive morphs, and the enigmatic evolution of exaggerated sexual traits in general.

---

The co-occurrence of highly different morphs within a single population is ubiquitous in nature with mimicry patterns in butterflies ^1,2^, floral forms in *Primula* ^3^, color morphs of stick insects ^4^ and alternative reproductive morphs in birds ^5–7^ as best-known examples. Development of these alternative phenotypes is generally under control of a single dominant locus that prevents the development of unfit recombinant phenotypes. Here, we investigate how selection shapes such a single locus determination mechanism that controls elaborate morphological differences involving multiple genes.

Two genetic models can explain the preservation and co-expression of adaptive gene combinations within each morph. First, genes involved in morph development could be controlled by a single regulatory factor that coordinates distinct genetic pathways ^8^. Second, co-adapted genes could be organized at the genomic level as a tight non-recombining cluster, referred to as a ‘supergene’ ^9,10^, inherited as a single genetic element. Multiple studies have shown that morph-determining alleles commonly differ by structural variations, such as inversions that suppress recombination along extensive multigenic chromosomal segments ^1,4–7^. It has been challenging to demonstrate that these structural variants effectively comprise a co-adapted gene combination. Low recombination in these regions indeed hinders fine-scaled mapping of individual gene effects. It is still largely unresolved whether structural variations at morph-determining alleles thus primarily affect a key-regulatory factor or suppress recombination among co-adapted genes. Moreover, both mechanisms could act synergistically if an initial structural mutation involving a morph-determining regulatory factor promotes subsequent linkage of genes subject to antagonistic selection between the morphs, ultimately leading to extensive chromosomal divergence, a mechanism also proposed for the evolution of sex-chromosomes ^11^.

The dwarf spider *Oedothorax gibbosus* is characterized by a male-limited genetic polymorphism. In this species, disruptive selection has led to the evolution of a “hunched” male morph (*f. gibbosus*) with conspicuous cephalic ornaments and an alternative “flat” morph (*f. tuberosus*) that lacks these features completely ^12^ (Fig. 1A,B, Fig. S1). The species belongs to the Erigoninae subfamily (dwarf spiders) where sexual selection has driven the evolution of bizarre and diverse male head structures ^13^ (Fig. S2). The cephalic region of the hunched morph of *O. gibbosus* consists of a large hump preceded by a groove covered with long innervated setae ^14^ (Fig. 1B). The structure functions as a nuptial feeding device and produces glandular secretions that allow hunched males to elicit copulations by previously inseminated females ^12,15^. Flat males lack these cephalic modifications completely and develop a female-like carapace (Fig. 1B). Hunched and flat males always co-exist in a stable equilibrium in which earlier maturity of flat males increases their access to unmated females early in the breeding season, while late developing hunched males have increased mating success with mated females ^12^.

**Figure 1.**
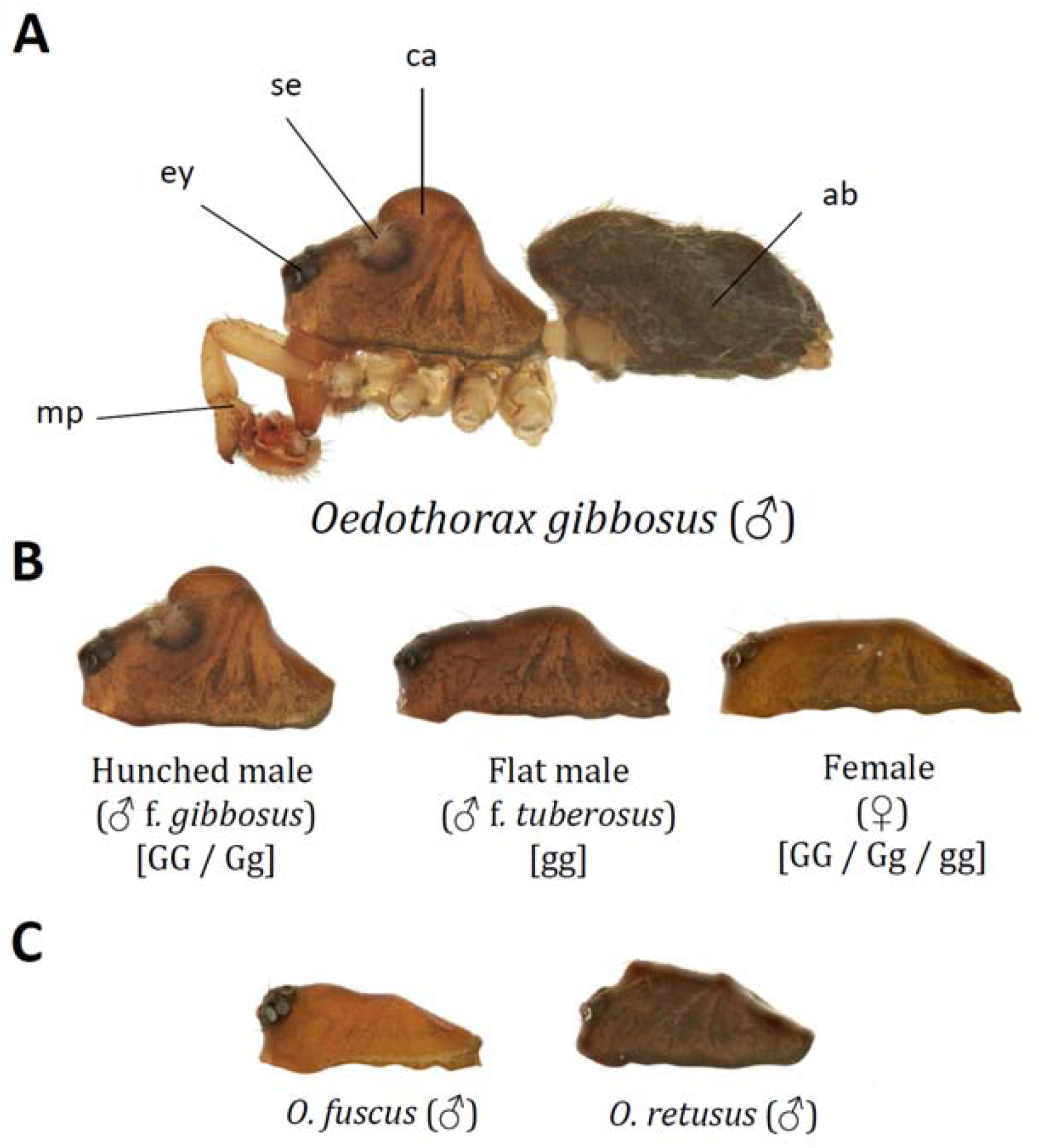
Head structures of *O. gibbosus* and outgroup species. **(A)** Habitus of a hunched male Oedothorax gibbosus f. gibbosus (legs are removed to visualize carapax)(ab = abdomen, ca = carapax, se = setae, ey = eyes, mp = male pedipalp). **(B)** Carapaces of a hunched male (*O. gibbosus* f. *gibbosus*, left), flat male (*O. gibbosus f. tuberosus, center*) and female *O. gibbosus* (*right*). *Genotypic morph determination is given between square brackets*. (*C*) *Male carapaces of the used outgroup species O. fuscus* (*left*) and *O. retusus* (*right*).

Although the complex phenotypic differences between hunched and flat males likely involve multiple genes, male development into one of these morphs is under control of a single autosomal locus (*G*-locus) with hunched (G) being dominant over flat (g) ^16^. In the present study, we unravel the genetic basis and evolution of these alternative phenotypes and show how duplications of male-specific genes, including a key regulator of sex-determination, resulted in a hunch-specific supergene that decouples the development of specific male traits from the highly conserved sex-determining system.

## Results and discussion

### Mapping of the *G*-locus

We first assembled the genome of *O. gibbosus* using PacBio reads originating from a pool of heterozygous hunched siblings and a pool of flat siblings that all originated from the same family (Tables S1, S2 and S3). The resulting genome sequence measures 821Mb (87% of the estimated genomes size; 7,804 contigs; N50 = 973kb) of which 54.4% was identified as repeat sequence (Table S3). Although this ‘heterozygote hunched assembly’ was used as our reference genome sequence, we also performed a separate ‘homozygous flat assembly’ using the reads of the pool of homozygous flat siblings only and used this to reconstruct the g-allele.

We mapped the G locus on a genetic map based on a set of 5,370 high-quality single nucleotide polymorphisms (SNPs) obtained from RADtag sequencing of offspring originating from a cross between a wild caught heterozygous (Gg) hunched male and homozygous gg female (Fig. S5). Associating both male offspring phenotypes with these SNPs localized the G locus (log_10_P > 3) at the distal end of the autosomal linkage group 3 (LG_3; Fig. 2A). Genotypes of SNPs with the strongest association (log_10_P = 7.8, 5.91cM on LG_3) segregated according to the expected genotypes at the *G* locus by being heterozygote in the hunched male parent and all hunched male offspring and homozygote in the female parent and all flat male offspring. These 87 SNPs were distributed over 22 contigs summing to a total length of 19.9Mb.

**Figure 2.**
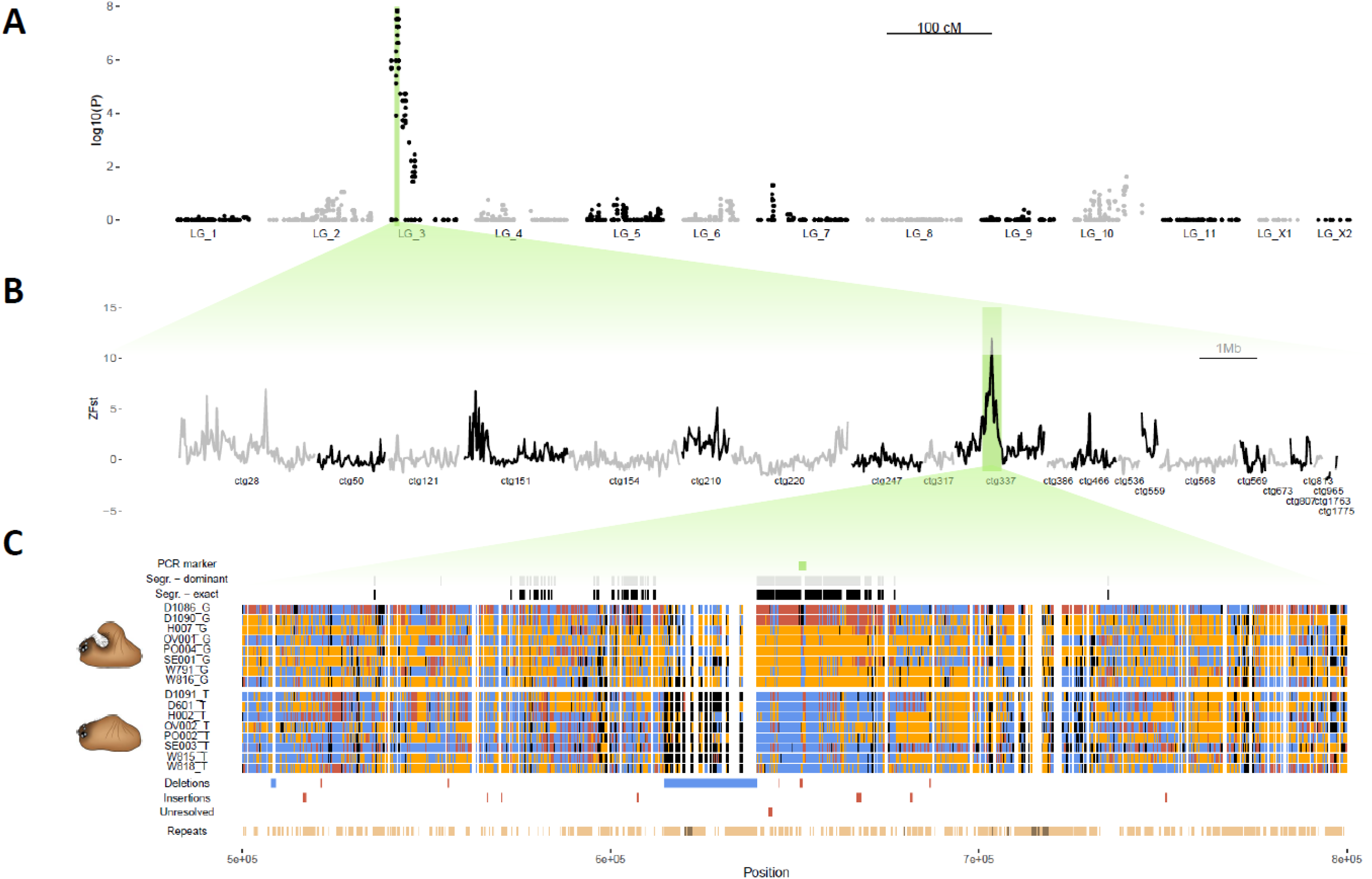
Genetic mapping, identification and characterization of the genomic region underlying the male reproductive morphs. **(A)** Association between SNP genotypes and male morph phenotypes on the *O. gibbosus* linkage map. **(B)** Genetic differentiation (ZF_ST_) between unrelated hunched (n = 8) and flat (n = 8) males at contigs identified by association analysis (20kb windows). (C) Detail of the most differentiated region in contig ctg337 (500kb – 800kb). Row 1 (PCR marker): location of the diagnostic PCR marker. Rows 2 and 3 (Segr. – dominant and Segr. – exact): location of SNPs with genotypes segregating according to dominance of hunched (grey) and according to the expected genotypes of the 16 resequenced individuals (black) respectively. Rows 4 – 19: SNP genotypes with color codes blue for the homozygote major allele, red for homozygote minor allele, yellow for heterozygote and black for missing genotypes. SNPs with minor allele frequencies < 0.1 were removed to emphasize morph specific genotype differences. Rows 20-22: Deletions (Deletions), insertions (Insertions) and insertions with unresolved length (Unresolved) as identified by Sniffles v 1.0.11^24^ based on PacBio read mapping. Red and blue bars indicate structural variations observed for the hunched and flat allele respectively. Row 23: Location of repetitive elements as identified by RepeatMasker v4.1.0 and RepeatScout v1.0.5 ^25^. Unclassified repetitive elements are coded light brown, LTR retrotransposons are coded dark brown.

Next, we resequenced the genomes of eight hunched (two GG and six Gg) and eight flat (gg) males, sampled in a pairwise manner, from six different populations and a single individual of the related species *O. retusus* and *O. fuscus* (Fig. 1C). Overall genetic differentiation among the 16 individuals was primarily structured by population origin and showed no differentiation according to male morph type (Fig. S6). In contrast, the genomic region identified in our linkage analysis showed an approximate 100kb region of elevated genetic differentiation (ZF_ST_) in the central part of contig ‘ctg337’ (Fig. 2B). This region was further characterized by an increase in sequence divergence (d_*xy*_) and high and low nucleotide diversity (π) in hunched and flat individuals, respectively (Fig. S7). SNPs at the center of this peak segregated according to the dominant expression of the hunched phenotype and were fully consistent with the expected genotypes at the *G*-locus of the sequenced males i.e., homozygous in the two GG males, heterozygous in Gg males and homozygous for the alternative allele in flat (gg) males (Fig. 2C). To further confirm the correct assignment of this genomic region in the expression of the alternative phenotypes, we screened an additional 88 adult males of different populations with a diagnostic marker spanning a ∼900bp deletion associated with the hunched-determining allele that is situated at the center of this region (Fig. 2C). This genotyping assay confirmed the presence of this deletion in all 35 phenotypic hunched males and in none of the 53 phenotypic flat males.

### Haplotype relationship at the *G*-locus

Haplotypes at the most strongly differentiated region (ctg337:640kb – 660kb) clustered into two highly divergent and non-recombining groups corresponding to the hunched- (G) and flat- (g) determining alleles (Fig. 3A). The coalescence time between these two alleles was the oldest compared to the maximum coalescence times observed in a random selection of 150 genomic windows of 20kb, which is consistent with long-term balancing selection maintaining two anciently diverged non-recombining alleles (Fig. 3B).

**Figure 3.**
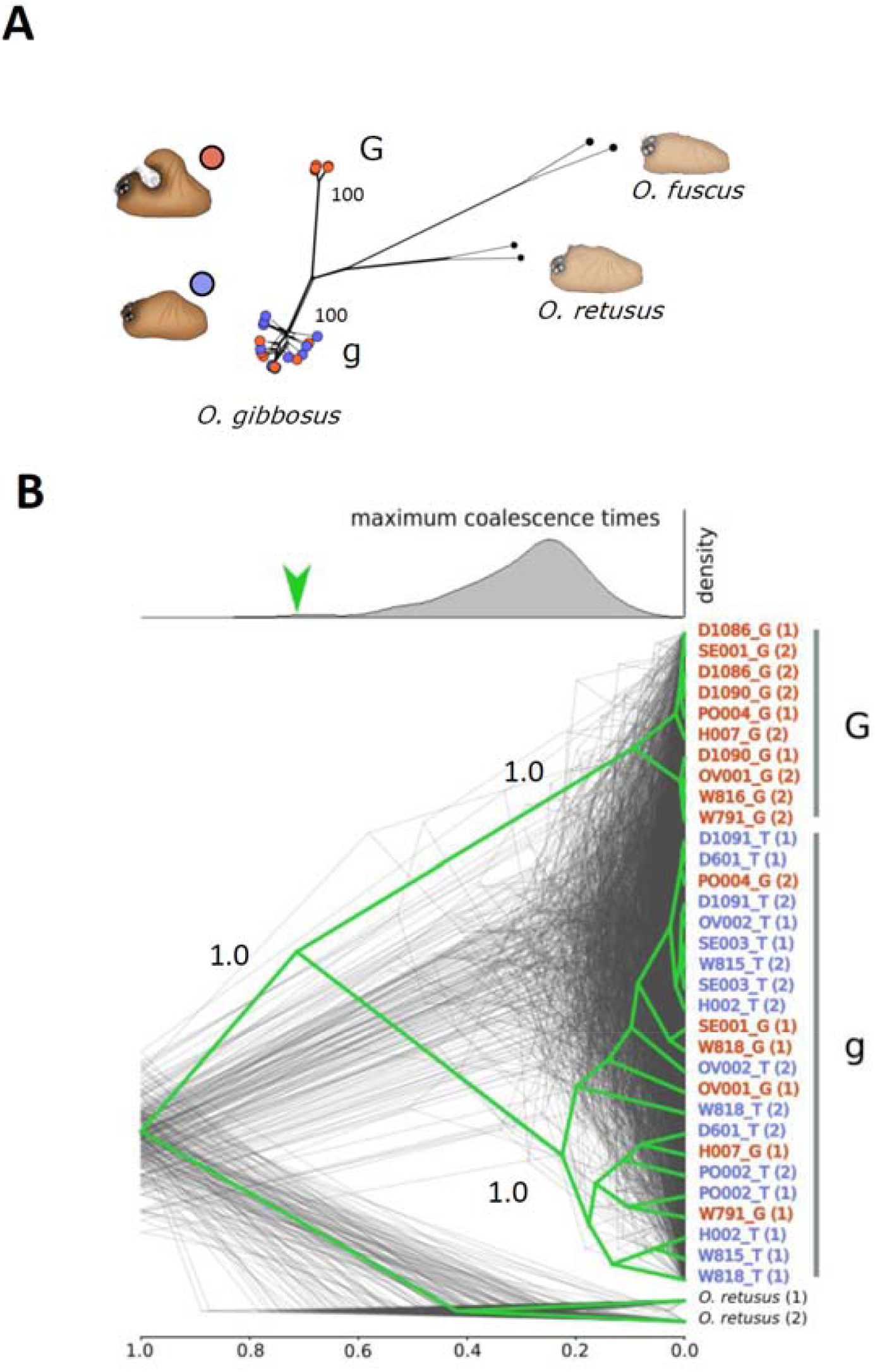
Haplotype relationships and recombination at the *G*-locus. (ctg337: 640kb – 660kb). (**A**) NeighborNet haplotype network at the *G*-locus with bootstrap support values of the two haplotype clades (**B**) Haplotype coalescence pattern at the *G*-locus (green, with branch labels depicting posterior clade probabilities) and across 150 randomly sampled 20kb windows (dark grey). Node ages are scaled relative to the divergence from the outgroup species *O. retusus*. Upper density plot depicts the distribution of the maximum coalescent times of the *O. gibbosus* haplotypes at the 150 randomly sampled 20kb windows, with the green arrow indicating the maximum coalescent time at the *G*-locus. Haplotypes originating from hunched and flat males are coded red and blue respectively in both graphs.

### An extensive insertion/deletion polymorphism distinguishes the G and g allele

Mapping long-read PacBio data originating from the heterozygous Gg (hunched) and homozygous gg (flat) offspring pools and the homozygous flat assembly identified the sequence at the *G*-locus in our heterozygote reference assembly as the g allele (Fig. S10). A structural variation (SV) analysis showed that the G allele structurally differs from this g allele by multiple insertions, including one of 25kb and one of unresolved length (Fig. 2C, Fig. S11). We therefore investigated if the G allele includes more extensive genomic fragments that are potentially represented as separate contigs in our heterozygote reference assembly. We detected these contigs by simultaneously testing each contig for higher coverage depth in hunched compared to flat individuals (t -test; P < 0.01) as well as the presence of positions that are consistently covered by short- and long reads in all hunched individuals, but in none of the flat individuals (Fig. S12). This selection procedure is unlikely to yield false positives as assessed by a random assignment of morph type across individuals (P < 0.01). Besides the contig containing the morph determining locus that we identified earlier (‘ctg337’), a total of 97 additional contigs matched these two criteria (Fig. 4A). Although these contigs were on average of small size (30.5kb), they summed to a total length of ∼3Mb. To confirm that no homologous, though potentially highly divergent, flat-specific contigs are present, we performed a more restricted contig depth comparison that includes the homozygote individuals only. If both alleles would comprise morph-specific sequences, we expect to find morph-specific contigs sets in either homozygote hunched or homozygote flat individuals. In contrast, we again observed this highly asymmetric coverage pattern with contigs showing higher coverage in hunched males only, but not in flat males (Fig. 4A, right panel). Lastly, we searched for a potential homologous flat sequence by mapping these 97 hunched-specific contigs to the remaining genome sequence. Although approximately one third of the total sequence length of these contigs (0.98Mb of the 3Mb) mapped with high quality to the remaining genome, it did not target a single genomic region but was spread across all linkage groups (Fig. 4B) and therefore likely represent mappings of repetitive or paralogous regions. Moreover, if the targets of this mapping would point towards a flat-specific sequence, we expect their normalized coverage to approximate zero in homozygote hunched and half in heterozygote hunched individuals compared to flat individuals (Fig. S13). In contrast, coverages at these mapping targets were highly similar for GG, Gg and gg individuals (Fig. S13) and are therefore unlikely to point towards a homologous g-allele. Based on these congruent findings from our haplotype assemblies, contig depth comparison and mapping approach we conclude that the main difference between both morph-determining alleles comprises an approximate 3Mb sequence fragment that is only present in the G allele. We further refer to this sequence as the ‘hunch-specific sequence’.

**Figure 4.**
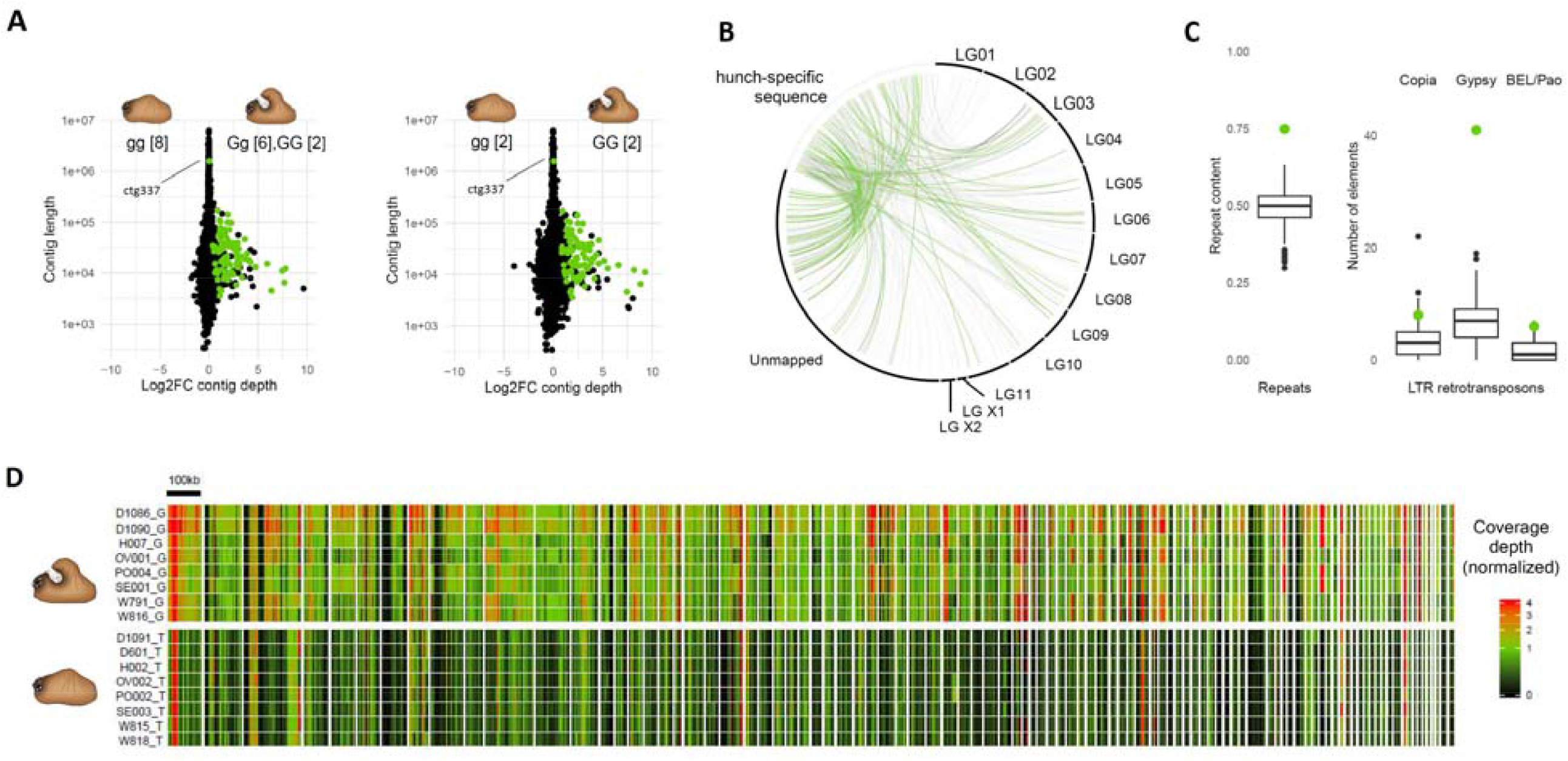
Identification and characterization of a hunch-specific sequence spanning ∼3 Mb. **(A)** Difference in contig coverage depth (log2 fold change) between the resequenced flat [8 homozygotes] and hunched [6 heterozygotes and 2 homozygotes] individuals (left panel) and for the 2 homozygote flat and 2 homozygote hunched individuals from population D only (right panel). Each dot represents a contig, with green dots representing contigs identified to be only present in hunched individuals (see text for details). (**B**) Mapping of the contigs of the unique hunch-specific sequence to the remainder of the *O. gibbosus* genome. Grey and green links depict mappings of non-coding and coding sequences, respectively. (**C**) Distribution of the repeat content (left) and density of LTR retrotransposons (right) across 3Mb sliding windows, with green dots depicting the observed density in the 3Mb unique sequence of the hunched allele. (D) Normalized coverage depth distribution (2kb windows) across the 97 contigs in the hunched allele identified as being deleted in the flat allele. Contigs are separated by blanks. Depth values higher than four were truncated.

### Gene and repeat content of the hunch-specific sequence

Coverage depth distribution across the hunch-specific sequence appeared highly uneven, peaking to values of more than 30 times the average genome coverage depth, indicating that it is largely composed of highly repetitive sequences (Fig. 4D). Repeat content analysis confirmed this by assigning 75% of the sequence as repetitive (Fig. 4C), which is substantially higher compared to the repeat content in similar sized regions of the genome (Fig. 4C). The sequence also contained exceptional densities of transposable elements (TE), particularly the LTR retrotransposons *Gypsy* and Pao and the non-LTR retrotransposon *R1/LOA* (Fig. 4C, Fig. S14). For the most abundant TE in the genome (*Gypsy*), the density of elements was about six times higher compared to the average number of elements found in similar sized regions of the genome, revealing that it constitutes among the most TE rich region of the entire genome (Fig. 4C, Fig. S14). This exceptional repeat content likely contributed to the difficulty to assemble the hunch-specific contigs as one contiguous sequence. We identified 193 predicted genes on the hunch-specific sequence, of which the majority (149 genes) showed clear, and often very high, similarities to genes located outside this fragment and thus present in the genomes of both morphs (Fig. 4B, Fig. S15, Table S4). Because these paralogous copies were distributed across the entire genome (Fig 4B), gene content of the hunch-specific sequence accumulated most likely by multiple duplication and translocation events rather than through a single segmental duplication.

Many of the genes located on the hunch-specific sequence likely lost their functionality as illustrated by the higher presence of internal stop-codons in their coding sequence (Table S4) and the significantly lower expression levels compared to their paralogs located in the remaining genome sequence (Fig. S16). For example, two-thirds (66%) of the genes located on the hunch-specific sequence showed no or only marginal (normalized expression cut-off < 1) expression compared to their paralogs located outside this fragment (10%, Fig. S16).

### Functional gene content identifies a Dmrt paralog on the hunch-specific sequence

A total of 40 genes showed clear hits to proteins with known functions (Table S4). One of these genes matched the *doublesex/mab3 related transcription factor* (Dmrt). Dmrt genes contain the highly conserved DM domain DNA binding motif that, though not exclusively, regulates the expression of sexually dimorphic traits in a very wide range of animal taxa like nematodes, insects and vertebrates, including humans ^17,18^. For many organisms, *Dmrt* genes act as masculinization genes, with knockdown inducing female development ^18^. If absence of this *Dmrt* gene in flat males is involved in the development of their female-like head structure, we predict its expression to be downregulated in genotypic hunched (Gg) females as well. RNA sequencing of genotypic hunched and flat males and females confirmed this prediction and revealed the clear association between the development of the complex cephalic structure and expression of this Dmrt gene (Fig. 5A). These findings suggest that the shared suppression of the elaborate male head structure in flat males and females is under control of the same gene, but that its downregulation involves very different mechanisms i.e., through a structural variation involving the absence of an entire gene (flat males) or through upstream regulatory pathways under control of the sex chromosomes (females).

**Figure 5.**
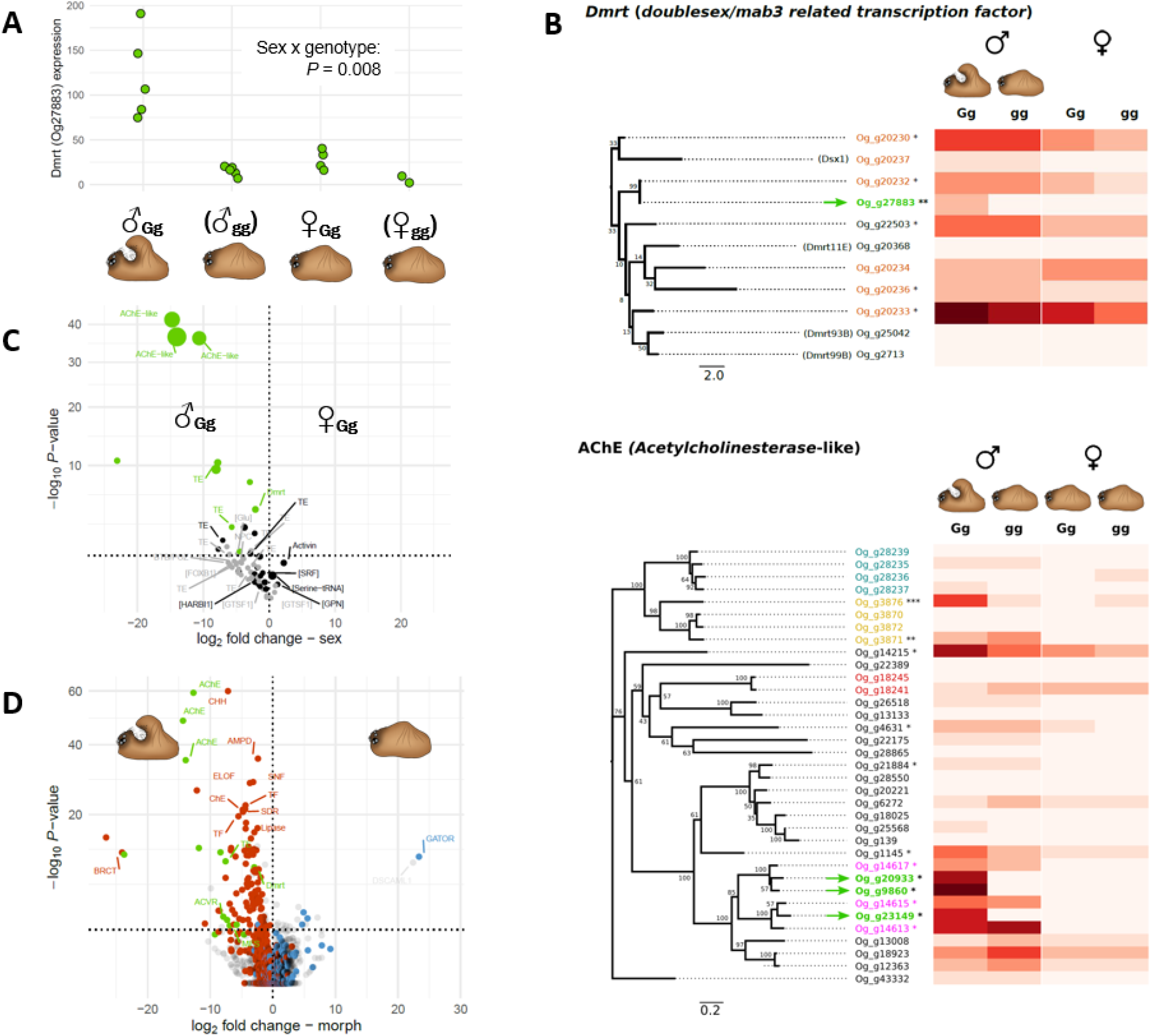
Morph- and sex-specific gene expression patterns. **(A)** Difference in expression between genotypic hunched (Gg) and flat (gg) males and females of the *doublesex/mab-3 related transcription factor* (*Dmrt*_Og27883) located on the hunch-specific sequence. Each dot represents the normalized individual read counts. Homozygote gg individuals, indicated with brackets, lack this gene and their expected expression level equals 0. (**B**) Maximum likelihood tree of the encoded protein sequences of the 11 *Dmrt* genes (upper panel) and 35 *Achetylcholinesterase*-like genes (lower panel) of *O. gibbosus*. The tree was constructed based on a COBALT-alignment with distances computed using the WAG substitution model and gamma distributed rate variation among sites. Bootstrap values are presented next to the branches. Genes clustered on the same contig are depicted with the same color code, with those indicated with a green arrow being located on the hunch-specific sequence. Heatmaps right of the trees show the corresponding expression levels in adult Gg and gg males and females, with darker values depicting higher expression values. Genes with an asterisk are significantly more expressed in males (*P*_*adj*_ < 0.05). (**C**) Sex-specific expression (log_2_ fold change) of all genes located on the hunch-specific sequence (Gg males versus Gg females). Dot size is proportional to the average normalized expression of each gene. Genes depicted in green are differentially expressed between the two male morphs and genes depicted in grey are only marginally expressed in adults. Genes located above the dotted horizontal line are significantly differentially expressed between the sexes (*P*_*adj*_ < 0.05). Genes with names between square brackets were partial fragments or showed premature stop codons. (**D**) Morph-specific gene expression (log_2_ fold change). Genes located above the dotted horizontal line are significantly differentially expressed between the male morphs (*P*_*adj*_ < 0.05). Genes coded in green are located on the hunch-specific sequence, genes coded in red are significantly higher expressed in Gg males compared to Gg females and genes coded in blue are significantly higher expressed in Gg females compared to Gg males.

Most metazoans contain multiple *Dmrt* genes, with some of them (Dmrt93B, Dmrt99B and Dmrt11E) likely controlling early developmental processes not related to sexual differentiation ^19^. For the *O. gibbosus genome*, we detected a total of 11 Dmrt genes, which is substantially higher compared to those in the well-annotated genome of the spider *Parasteatoda tepidariorum* (7 genes)^20^ and among the highest ever recorded in a metazoan ^19^. Six of these appeared to be physically clustered within a ∼86kb genomic region that mapped adjacent to the *G*-locus in our linkage map (ctg151:154kb – 240kb; Fig. 1B). These *Dmrt* genes were structurally highly divergent and five did not show any clear phylogenetic relationship with published *Dmrt* sequences of the spider *P. tepidariorum* (Fig. S17). The *Dmrt* gene located on the hunch-specific sequence (Og_g27883) was strongly related to one of these physically linked *Dmrt* genes (Og_g20232) and suggests that it arose from a duplication event (Fig. 5B, Fig. S17). Four of the genes in this cluster, including the *paralogous Dmrt* gene (Og_g20232), were significantly upregulated in both hunched and flat males compared to females, which indicates their shared developmental role in sexual differentiation (Fig. 5B).

We further identified three *acetylcholinesterase*-like (*AChE*-like) genes on the hunch-specific sequence, which were among the 1% most highly expressed genes in adult hunched males. Their expression pattern and evolutionary history showed remarkable similarities to those of *Dmrt* (Fig. 5B). First, expression was completely suppressed in females as inferred from genotypic Gg females, being carriers of these three *AChE*-like copies (Fig. 5B). Second, their phylogenetic relationship with other *AChE*-related genes of *O. gibbosus* reveals that these three genes also likely originated from a recent duplication from a physical cluster of related *AChE*-like genes present in the genomes of both morphs (Fig. 5B, Fig. S18). Third, these paralogous AChE-like genes located in the genomes of both morphs were also strongly downregulated in females (Fig. 5B). The strong male-biased expression of these *AChE*-like genes as well as their paralogs, all being among the top 0.1% most differentially expressed genes between both sexes, also point towards a highly sex-specific functional role. AChE genes are best known for their hydrolysis of the neurotransmitter acetylcholine at neural synapses, a function that is not directly expected to relate to sexual differentiation. In contrast to vertebrates, however, invertebrates and in particular arachnids possess multiple AChE-related genes with non-classical and non-neuronal functions ^21,22^. Yet, their involvement in sexual and morph differentiation in *O. gibbosus* and spiders in general remains unclear.

Of the remaining functionally annotated genes located on the hunch-specific sequence, we identified two additional genes encoding for transcription factors involved in developmental processes i.e. Forkhead box protein and *BTB/POZ* domain-containing protein, but these were not expressed in adults. Lastly, eighteen genes were related to (retro-)transposable elements (*Gag-Pol polyprotein, DNA transposase*) and 13 genes constituted partial fragments or contained premature stop-codons and were considered as pseudogenes that lost functionality.

### The hunch-specific sequence is enriched with male-expressed genes

The male-specific expression of *Dmrt* and *AChE* and their paralogs suggest that the functional gene content of the hunch-specific sequence consist primarily of duplications of male-specific genes. We tested this more generally by comparing the sex-specific expression of all genes located on the hunch-specific sequence (both functionally annotated as well as non-annotated) between Gg males and Gg females. Of the eleven genes located on the hunch-specific sequence with evident expression in hunched males, ten were significantly downregulated in Gg females (P_adj_ < 0.05; Fig. 5C). Average and median expression of these genes was also significantly more male-biased compared to those of a similar sized random sample of genes in the remaining genome (Fig. S19, randomization test, P < 0.0001) and primarily caused by a lack of female expression (Fig. S19). The closest paralogs of these genes, which are located outside the hunch-specific sequence and thus present in both morphs, show a likewise male-biased expression (Fig. S20; r_P_ = 0.53, P < 0.0001) and were also on average significantly more male-biased expressed compared to a random sample of genes in the genome (randomization test, P < 0.0001; Fig. S19). Thus, functional gene content of the hunch-specific sequence appears to have expanded by duplications of male-expressed genes.

### Upregulation of male-specific genes in hunched males

Presence of a male-specific transcription factor (*Dmrt*) suggests that the hunch-specific sequence upregulates additional male-specific genes in hunched males. Expression profiling of both male morphs across all genes confirmed this by showing a highly asymmetric pattern with 125 genes being upregulated in hunched compared to flat males, while only 21 genes were upregulated in flat compared to hunched males (Fig. 5D, Fig. S21, Table S5). Most of the genes upregulated in hunched males (87%) were significantly downregulated in females as well, which suggests that these genes are either male-specific or associated with the development of the hunch. To distinguish between these two alternatives, we further tested if genes upregulated in hunched compared to flat males are also upregulated in flat males compared to females as both are devoid of hunch development. This comparison revealed that of the 125 genes upregulated in hunched compared to flat males, 31 are also upregulated in flat males compared to females (Fig. S21). Thus, transcriptomic differences between both male morphs are not restricted to hunch development, but additionally comprise upregulation of general male-specific genes in hunched males.

### Evolutionary history of the supergene

The extensive divergence between both alleles could either have been triggered by an initial duplication of a morph-determining factor, with Dmrt being the most likely candidate, which resulted in an incipient G-allele, or through the deletion of this duplicated gene resulting in the g allele. We tested for the presence of the hunch-specific sequence in the two outgroup species *O. retusus* and *O. fuscus*, which revealed that coding sequences located on the hunch-specific sequence are significantly less covered in these two outgroup species (P< 0.001; Fig. S22A). For *O. retusus* and *O. fuscus*, coverage depths were like those in flat males that lack these duplicated genes (Fig. S22A) and suggests that the duplicated genes located on the hunch-specific sequence originated after the species split from *O. retusus*. A detailed phylogenetic analysis on the Dmrt gene confirmed this by placing the duplication event of this gene more recently compared to the divergence between *O. gibbosus* and *O. retusus* (Fig. S22B). The estimated timing of this duplication coincided with the estimated timing of the divergence between the G and g allele (Fig. S22B), which at first suggests that this duplication event initiated this polymorphism. However, the large credibility intervals of these divergence time intervals do not allow to infer the order of these two events and, hence, to discriminate if the Dmrt duplication or its secondary deletion initiated the divergence between both alleles.

## Conclusion

Secondary sexual traits and their intraspecific polymorphisms represent one of the most astonishingly diverse and spectacular morphological variations in nature, but their evolution is still surprisingly poorly understood ^23^. We here show how a large indel polymorphism, comprising a morph-specific multigenic sequence fragment, controls a complex male dimorphism. The functional gene content of this fragment constitutes a duplicated key regulatory gene in sexual differentiation (*Dmrt*) as well as additional duplications of male-specific genes (Fig. 6). These findings demonstrate that the hunch-specific sequence acts as a masculinizing supergene that underlies the development of the exaggerated secondary sexual traits in hunched males. The unique male expression of the genes on this fragment further resolve how expression of this dimorphism is limited to one sex.

**Figure 6.**
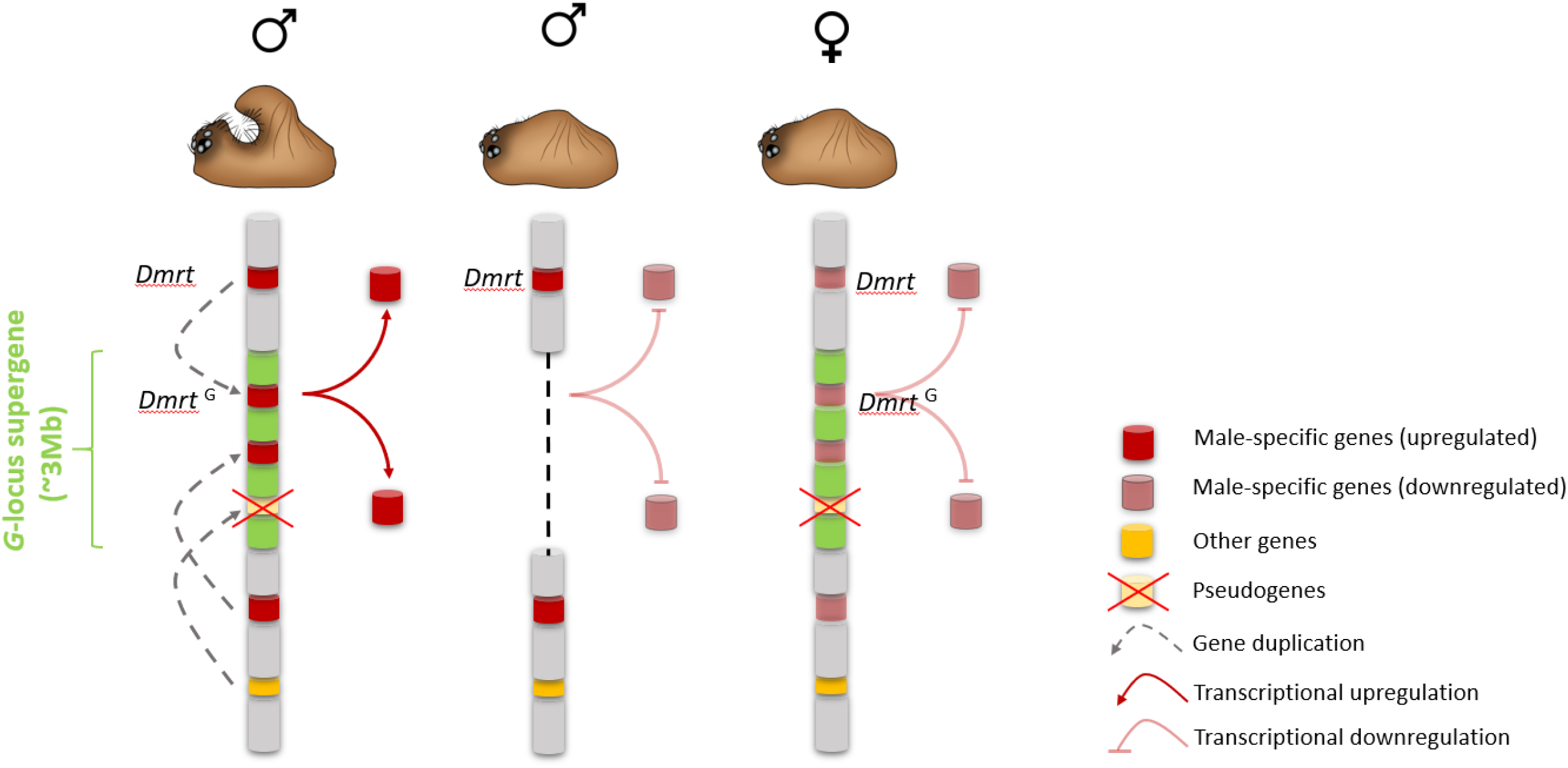
Schematic presentation of the proposed structure and function of the morph determining alleles at the *G*-locus in males and females. The hunch-determining allele comprises a hunch-specific sequence (green), which is absent in the recessive flat-determining allele. The fragment contains duplicates of male-expressed genes (red), including regulatory elements (*Dmrt*) that control the downstream upregulation of additional male-specific genes. In females, male-specific genes like those located at the supergene are downregulated.

While chromosomal inversions, and concomitant suppressed recombination, are well known drivers of supergene evolution ^9,10^, results from our study indicate that extensive indel polymorphisms may be a yet underappreciated mechanism controlling morph differentiation. Moreover, the genome-wide distribution of genes paralogous to those within the hunch-specific sequence strongly suggests that this morph-specific fragment evolved through an initial structural variation involving a morph-determining factor and subsequent recruitment of genes positively selected in hunched males only. Evolution of supergenes by accumulation of “gain of function” genes for one single morph not only explains how deleterious effects are prevented in homozygous individuals that lack such multigenic fragments, but further provides unique support to the debate that the structural differences between morph-determining alleles primarily emerge because of reduced recombination, rather than causing it ^10^.

## Acknowledgments

We thank A. Mariscal for help in RNA extraction; K. Smistek and C. Locatelli for help in taking pictures of the specimens; F. De Block, S. Cogneau, V. Vandomme and E. Veltjen for help in sampling and breeding; L. Sterck for advice on genome annotation and gene curation; N. Edelman for sharing code to depict phylogentic trees; M. Horn and T. Halter (University of Vienna) for help in the identification of contaminant bacterial contigs. The computational resources (Stevin Supercomputer Infrastructure) and services used in this work were provided by the VSC (Flemish Supercomputer Center), funded by Ghent University, FWO and the Flemish Government – department EWI.

## Author Contributions

FH conceived the project. FH, GS and CV performed the experiments. FH, ZDC, SMVB, SK and CV analyzed the data. FH drafted the manuscript with input from all authors.

## Data and materials availability

DNA sequences have been deposited in the NCBI SRA (PRJNA681589).

## Material and Methods

### Genome assembly

We generated multiple *de novo* genome assemblies based on three independent library preparation methods (Table S1) and selected the version with the highest contiguity and completeness statistics as final genome assembly for downstream analyses. A short-read assembly (**“Ogibo_platanus”**) was constructed using Illumina reads (HiSeq2000) from three paired-end libraries (2x 100bp) (insert sizes of 170bp, 500bp and 800bp) and two mate-paired libraries (2×50bp)(insert sizes of 2kb and 5kb) and was assembled with the Platanus assembler ^26^. Libraries were constructed from DNA extracts obtained from offspring of a cross between a heterozygous Gg male (D1077) and a gg homozygous female (D1076) from population D. The second assembly (**“Ogibo_10x”**) was generated based on linked reads produced with 10x Genomics Chromium chemistry originating from a single heterozygous hunched male (PO11_22) and assembled with the Supernova assembler (v2.1.1., 10x Genomics). Lastly, two separate 20kb PacBio SMRTBell libraries were constructed from a pooled sample of 12 heterozygous (Gg) individuals and a pooled sample of 6 homozygous flat (gg) individuals. All these individuals originated from the same family involving a cross between a homozygous flat (gg) male (W776) and a heterozygous (Gg) female (W744). Each pooled sample was sequenced with 3 SMRT cells (6 SMRT cells in total). Sequences from both pools were assembled with Canu v.1.8 ^27^ (**“Ogibo_pacbio_canu”**) and wtdbg2 ^28^. Assembly with Wtdbg2 was performed based on the reads of both pools, resulting in a heterozygote assembly (**“Ogibo_pacbio_wtdbg2_het”**) as well as the pool of flat offspring only, resulting in a homozygote flat assembly (**“Ogibo_pacbio_wtdbg2_flat”**)

### Short-read assembly

Total DNA was extracted (NucleoSpin® Tissue kit, Macherey-Nagel GmBH) from 51 offspring (6 *gibbosus* males, 6 *tuberosus* males and 39 females) originating from a single cross between a heterozygous Gg *gibbosus* male (D1077) and a female (D1076) that were captured as immatures in population Damvallei (Table S1). Illumina paired-end (100 bp) and mate-paired (50 bp) libraries with insert sizes of 170 bp, 500 bp, 800 bp, 2 kb and 5 kb were constructed and sequenced at the Bejing Genomic Institute (BGI, Hongkong) on an Illumina HiSeq2000 system according to the manufacturer’s protocol (Table S1). Sequencing of the libraries resulted in a total of ∼72.2 Gb of raw sequencing data. Raw reads were corrected for sequencing error with SOAPec v2.02 ^29^, using a *k*-mer size of 17 and a low frequency cutoff of consecutive *k*-mer of 11. Frequency distribution of the k-mer species yielded an estimated effective sequencing depth of 44.4 x and a corresponding estimated genome size of 948Mb. We used Platanus v1.2.1 ^26^ to generate a genome assembly based on the error-corrected short-reads. Contigs were assembled using the 170bp, 500bp and 800bp libraries only. Both paired-end and mate-paired (2kb, 5kb) libraries were then used to scaffold the contigs (*platanus scaffold*) and to fill the remaining gaps connecting the contigs (*platanus gap_close*). The assembly (**“Ogibo_platanus”**) resulted in 589,217 scaffolds (85,264 scaffolds with length > 500bp), summing to a total length of 811Mb, and an N50 of 78,244bp (Table S2).

### Chromium 10x Genomics assembly

Total DNA was extracted from a hunched male (PO11_22) originating from a cross between a flat male (gg)(PO07) and a female of unknown genotype (PO11), which were captured as immatures from population Pollismolen. The emergence of both flat (n = 7) and hunched (n = 10) male offspring in this cross allowed us to infer the genotype of the extracted hunched individual as being heterozygous (Gg). HMW was extracted using the “DNA extractions from single insects” protocol provided by 10x Genomics® (v. 01/12/2018). Fragment analysis of the extracted DNA revealed that most fragments ranged in size between 6kb and 100kb, with a peak at 39kb. A sequencing library of barcoded linked reads was prepared at the Leiden Genome Technology Center (LGTC, The Netherlands) using Chromium Genome Library & Gel BeadKit v.2 (10X Genomics), Chromium Genome Chip Kit v.2 (10X Genomics), Chromium i7 Multiplex Kit (10X Genomics) and Chromium controller according to the instructions provided by the manufacturer. The library was sequenced (2×150bp) on a single cell of the Illumina NovaSeq platform resulting in 473M raw reads (∼75x coverage) (Leiden Genome Technology Center, The Netherlands)(Table S1). We used Supernova (v2.1.1., 10x Genomics) with default parameters to trim the reads and assemble the phased genome. An effective coverage of 50x was retailed after raw read processing. *K*-mer analysis by Supernova estimated a genome size of 939Mb. We specified the --style=pseudohap parameter to output a phased haplotyped genome assembly. The final assembly resulted in 47,583 scaffolds (5,690 > 10kb) with a total length of 873Mb and an N50 of 564kb (**“Ogibo_10x”**, Table S2).

### PacBio assembly

DNA of 6 homozygous flat (gg) and 12 heterozygous hunched (Gg) offspring, all originating from a single cross between a homozygous flat (gg) male (W776) and a heterozygous (Gg) female (W744), was extracted individually with the MagAttract® HMW DNA extraction kit (Qiagen) following the manufacturers protocol. We generated two pools of DNA extracts with one pool including all DNA extracts of the homozygous flat (gg) offspring and a second pool including the DNA extracts of the hunched (Gg) offspring. A 20kb SMRTbell template library was constructed for each pool separately and sequenced individually on the PacBio Sequel system using 3 PacBio SMRT® sequencing cells per library (6 PacBio SMRT® sequencing cells in total)(Macrogen Europe). This resulted in a total of 38.6 Gb (4.10M subreads) and 43.8Gb (4.75M subreads) output for the hunched and flat offspring pool respectively (Table S1).

Raw uncorrected reads of the hunched (Gg) offspring pool were corrected, trimmed and assembled with Canu v.1.8 ^27^ using default parameter values for the metaparameters “rawErrorRate (0.3)” and “correctedErrorRate (0.045)”. We further specified a genome size of 1Gb and discarded reads with a minimum read length of 1kb in all steps. The final Canu assembly (**“Ogibo_pacbio_canu”**) consisted of 39,205 contigs with a total length of 1.37Gb and an N50 of 48kb (Table S2). The contiguity of the assembly was further improved by scaffolding contigs with the Chromium 10x linked reads using the ARKS/LINKS scaffolding pipeline ^30,31^ with recommended parameter settings: -c 5 -j 0.5 -o 0 -m 50-10000 -k 30 -r 0.05 -e 30000 -z 500 -d 0 (ARKS) and - x 1 -l 5 -a 0.3 -z 500 (LINKS). This scaffolding step slightly decreased the number of scaffolds from 39,205 to 36,593 and increased the N50 from 48,106 to 57,828bp (**“Ogibo_pacbio_canu+links”**, Table S2).

A second PacBio assembly, based on the raw uncorrected PacBio long-reads of the hunched (Gg) and flat (gg) offspring combined, was generated with Wtdbg2 ^28^ using default parameter settings and assuming a genome size of 1Gb (**“Ogibo_pacbio_wtdbg2_het”**). The derived consensus sequences were initially subjected to a polishing step based on the original raw PacBio reads with the wtpoa-cns consenser included in Wtdgb2. We then performed two subsequent rounds of polishing based on short-read resequencing data (∼35x coverage) of a Gg heterozygote male (W791_G) originating from the same population (Table S1). The first polishing was conducted with the wtpoa-cns consenser and the second polishing round was conducted with HyPo v.1.03 ^32^. The wtdgb2 assembly consisted of 8,072 contigs with a total length of 833Mb and an N50 of 968kb.

We also generated a Wtdbg2 assembly using the same assembly procedure but including only the data of the pool of homozygous flat individuals (**“Ogibo_pacbio_wtdbg2_flat”**). Polishing was conducted similarly to the heterozygote assembly but based on the reads of the flat individuals only and short-read resequencing data (∼35x coverage) of a homozygote flat (gg) male (W815_T) originating from the same population (Table S1). This homozygous flat assembly consisted of 6,754 scaffolds (N50 = 525kb) summing to a total of 774Mb.

### Final genome assembly

Highest assembly contiguity was achieved from the wtdbg2 assembly based on PacBio reads originating from both the hunched and flat offspring pool (**“Ogibo_pacbio_wtdbg2_het”**, Table S2). This assembly also outperformed the other assemblies in terms of completeness, as assessed on a set of 1066 benchmarked universal single-copy orthologs (BUSCO) in arthropods ^33^, which recovered 88 % single-copy [C], 3.8% duplicated [D] and 3.0 % fragmented[F] orthologs (Table S2) and was used as final genome assembly (Table S3).

### Removal of contaminant scaffolds

Illumina reads were initially *de novo* assembled with SPAdes v3.9.1 ^34^ with default settings and the “--meta” parameter for metagenomic samples. Subsequently, binning of metagenomic assembled genomes (MAGs) of four known bacterial symbionts of *O. gibbosus* including *Rickettsia, Wolbachia, Rhabdochlamydia*, and *Cardinium* species ^35^ was performed with the mmgenome workflow ^36^ (https://github.com/MadsAlbertsen/mmgenome). Targeted re-assembly of the four bacterial MAGs was carried out using BBmap v32.14 (https://sourceforge.net/projects/bbmap/) and SPades v3.9.1 as described in ^37^, and MAG quality was assessed with CheckM ^38^. We next mapped PacBio reads to the symbiont MAGs with minimap v2.17 ^39^ and then *de novo* assembled the mapped reads with Unicycler v0.4.8 ^40^ with default parameters. The quality of the final symbiont MAGs was again assessed with CheckM ^38^. For the identification of contaminant scaffolds in the host genome assembly we taxonomically classified contigs with Kraken v2.0.7 ^41^. We first initialized a database using *kraken2-build* including human, viral, plasmid, protist, fungi, plant, and bacterial genome sequence databases, and then added the four symbiont MAGs with “--add-to-library”. Using *kraken2* with “--confidence 0.5”, we identified in total 254 putatively contaminating scaffolds (11.78 Mb) from multiple origins. The final assembly consists of 7924 contigs summing to a total size of 825Mb (87% of the k-mer estimated genome size of 948Mb), an N50 contig size of 973kb.

### Linkage map construction

Contigs of the reference genome were positioned on a linkage map that was obtained from a cross between a hunched male (W834) and a female (W828) that were sampled as juveniles at population W. Both individuals were raised individually till adulthood in the lab and mated. Spiderlings were isolated three days after their emergence from the egg cocoon and raised individually till adulthood. The cross resulted in 14 hunched, 13 flat and 19 female offspring. The proportions of hunched and flat offspring allowed us to infer the parental genotypes as being Gg heterozygote for the male parent and gg homozygote for the female parent (*P*_(female = Gg)_ = 0.0048; Exact binomial test).

Parents and all 46 offspring were genotyped by means of Restriction-site Associated DNA (RAD) sequencing based on an *Sbf*I restriction digest following the protocol described in ^42,43^ (Table S1). Libraries were sequenced on a single lane of an Illumina HiSeq1500 sequencer platform (Center of Medical Genetics, UA, Antwerp, Belgium). Assessing the presence of the restriction site, demultiplexing and quality filtering of reads was performed with the *process_radtags* tool in Stacks v.1.20 ^44^. Reads with identical randomly sheared paired-end sequences were considered as PCR duplicates and filtered with the *clone_filter* tool in Stacks. Filtered reads were mapped to the reference genome (“Ogibo_pacbio_wtdbg2_het”) with BWA (bwa-mem) producing individual bam files. SNPs were subsequently called with the UnifiedGenotyper walker in the Genome Analysis Toolkit (GATK)^45^. We generated a high-quality SNP set from the resulting VCF-file with VCFtools^46^ by keeping only biallelic SNPs with a genotype quality > 30 that are present in at least 42 individuals. Distribution of the average depth of the 9179 retained SNPs revealed depth peaks at ∼75x and ∼30x, corresponding to SNPs called in the forward and reverse reads respectively, and a sharp drop in the number of SNPs with depths exceeding 90x (Fig. S3). SNPs with depths above 110x (452 SNPs) were excluded as these might potentially include SNPs within repeat regions.

We used Lep-MAP2 ^47^ and associated scripts for the construction of the linkage map. We first obtained the posterior probabilities of the individual genotypes for the filtered SNP set with the script “pileup2posterior.awk” based on the samtools mpileup output. This step included an additional filtering step that excluded reads with a lower mapping quality (-q 60). The ParentCall module was used to call parental genotypes and to identify markers located on the X-chromosomes using a logarithm of odds (LOD) = 2. Markers showing high distortion from a mendelian segregation pattern were removed by the Filter module using the default dataTolerance of 0.01. The final set consisted of 5,369 high quality SNPs for linkage map construction.

The karyotype of *O. gibbosus* is 2n = 22,2X_1_X_2_ for females and 2n = 22, X_1_X_2_0 for males and corresponds to the general pattern of Entelegyne spiders (J. Král, pers. comm.). This is expected to result in 11 linkage groups corresponding to the 11 autosomal chromosomes and two linkage groups with X-linked markers corresponding to the two X chromosomes. Markers were assigned to linkage groups with the SeparateChromosomes module using LOD score limits ranging from 3 to 8. A LOD score limit of 7 resulted in 12 linkage groups with > 100 markers (Fig. S4). Location of the markers on the reference genome revealed that all markers assigned to LG12 were located on RADtags or scaffolds whose remaining markers were assigned to LG11, suggesting that LG11 and LG12 refer to the same chromosome. Inspection of the parental genotypes of markers on these two LG’s revealed that all markers on LG11 were heterozygote in the female parent and homozygote in the male parent, while a reverse pattern was observed for markers assigned to LG12 and thus split due to an absence of markers informative in both the male and female parent. As LG12 did not include unique scaffolds, we only considered LG11 in the linkage map. The largest linkage groups including the sex-linked markers were assigned to X1 and X2.

Markers within linkage groups were subsequently ordered by the OrderMarkers module, wherein we iteratively removed markers with a genotype error rate estimate > 0.1 ^47^. Ten independent runs per LG were performed and kept the marker order with the highest likelihood. The final map included 4,095 SNPs, distributed over 577 scaffolds constituting a total length of 543 Mb (65% of the assembled genome size)(Fig. S5).

### Genetic mapping of the G locus

We mapped the G locus on the linkage map with a parametric linkage analysis by calculating the two-point logarithm of odds (LOD) scores with the R package *paramlink* ^48^. The analysis was restricted to the male offspring as females do not express the distinct phenotypes. In accordance with the mendelian autosomal dominant inheritance mode of *gibbosus* as inferred from previous breeding experiments ^16^, the locus was specified as autosomal dominant with full penetrance (setModel(x,model=1)) and a frequency equal to 0.35 in our pedigree (dfreq=0.35). LOD scores were calculated for all 5,370 SNPs for a recombination rate between *G*-locus and SNP-marker genotypes equal to zero (theta=0).

### Individual sampling and breeding of resequenced individuals

We resequenced the genomes of eight flat and eight hunched males that were sampled pairwise in six different populations, with an average distance of 65 km, in Belgium. A single adult male of each morph was sampled in the populations Honegem (H: 50.9547°N, 3.9956°W), Overmere (OV: 51.0517°N, 3.9839°W), Pollismolen (PO: 51.1150°N, 5.6431°W) and Sevendonck (SE: 51.2757°N, 4.9368°W), while two males of each morph were sampled at population Walenbos (W: 50.9242°N, 4.8649°W). Field captured hunched males are most likely heterozygous Gg as the estimated frequency of the G allele in wild populations is approximately 0.2 ^12^, corresponding to 11% of the hunched males being homozygous GG. To assure that also homozygous (GG) hunched males were included in our selection of 16 males, we performed a breeding experiment with individuals captured at population Damvallei (D: 51.0570°N, 3.8302°W) wherein wild caught hunched and flat males were paired with females captured as immatures in the field. We separated offspring families wherein pairing with a flat male resulted in at least 10 flat male offspring only (F1 ‘flat families’ consisting exclusively of gg offspring) and families wherein matings with a hunched male resulted in a high proportion (∼75%) of hunched offspring (F1 ‘hunched families’ enriched for the G allele), which likely contain at least 25% GG males. To identify F1 GG males, we mated them with females of the pure flat families and males that produced at least 10 hunched and no flat male offspring were considered homozygous GG. This breeding experiment allowed us to isolate two homozygous GG males from population D that were not of the same family. We also included two field captured flat males from this population to maintain our pairwise sampling design. In addition, we resequenced a single male of the related species *Oedothorax retusus* and *Oedothorax fuscus*, which were used as outgroup.

### Whole genome resequencing, genotyping and sequence divergence

DNA of the 18 resequenced specimens was extracted with the NucleoSpin Tissue Kit (Macherey-Nagel GmBH) and TruSeqNano gDNA libraries were sequenced on an Illumina HiSeqX platform (Macrogen) resulting in between 20.0 Gb and 39.6 Gb, corresponding to ∼21x - ∼41x coverage per specimen (Table S1). Reads were mapped with BWA v0.7.17 (bwa mem)^49^ to the Ogibo_pacbio_wtdbg2_het assembly and reads with a minimum mapping quality < 30 were filtered with SAMtools ^50^ (-q 30). SNPs were called with the UnifiedGenotyper tool implemented in the Genome Analysis Tool Kit (GATK) v3.8-0 ^45^ using the default read filter settings. Only individual genotypes of biallelic SNPs that are present in all individuals with a genotype quality higher than 30 were retained using VCFtools v0.1.16 ^46^ (--minGQ 30 --max-missing 1 --min-alleles 2 -- max-alleles 2 –maf 0.01 –max-maf 0.99), resulting in a total of 29.570.023 high quality SNPs. The same filtering procedure was applied excluding the outgroups species *O. retusus* and *O. fuscus* and resulted in 19.541.199 SNPs for the *O. gibbosus* individuals only. Differentiation (Fst) between the two morph types was calculated based on non-overlapping 20kb windows with VCFtools^46^. Sequence divergence (d_XY_) and nucleotide diversity (π) were calculated in non-overlapping 10kb windows with the Egglib package ^51^. SNPs with genotypes that segregated according to the expected genotypes of the resequenced individuals were identified based on an *in-house* script (“GetGibPos.py”) that searches the VCF file for positions with homozygous genotypes in the two lab bred homozygous hunched individuals from population D, heterozygous genotypes in the remaining field capture hunched males and homozygous for the alternative allele in all flat individuals.

### PCR genotyping at the *G*-locus

We developed a diagnostic marker around a 900bp deletion in the hunched allele. This deletion is positioned within the genomic region that showed the strongest differentiation between the two male morphs (Fig. 2, ctg337: 651,675kb – 652,592kb). The primers Gib_F (5’- CGAGGCGTTGGTTTGAAAGG) and Gib_R (5’ – AGCTTGGAAGGGAAGCAAGA) were designed using Primer3 v. 2.3.7 ^52^ and anneal at both sides of the 900bp indel resulting in amplicons of ∼760bp (G allele) and ∼1660bp (g allele) that can easily be distinguished on a gel. PCR’s were conducted for 40 cycles with the Phusion^™^ High-Fidelity DNA polymerase (NEB) using an annealing temperature of 65°C.

### Genetic structure, phasing and phylogenetic relationships

Genetic structure among the resequenced individuals was investigated by a Principal Coordinate Analysis (PCoA) implemented in the adegenet v 2.1.2 ^53^ package in R v3.5.3. High quality SNPs present in all individuals were first thinned to 1 SNP per 20kb, resulting in total of 39,366 SNPs used for the PCoA.

We reconstructed haplotype relationships at both the G locus (ctg337:640kb-660kb) as well as a random set of 150 windows of 20kb sampled across the genome. Resequencing data of the eight unrelated flat, eight unrelated hunched and the outgroup species *O. fuscus* and *O. retusus* were used for this analysis. Windows were selected based on a random selection of 150 positions in the genome, followed by the extraction of SNPs positioned 20kb downward on the respective contig. SNPs at each window were phased with SHAPEIT2 ^54^ using both read information and population information. Phase informative paired reads were first extracted with the extractPIRs tool and subsequently used together with population information to reconstruct haplotypes. SplitsTree v4.14.4 ^55^ was used to generate a NeigborNet haplotype network based on the uncorrected p-distances at the *G* locus. Haplotype trees at all windows were reconstructed by running a multispecies coalescent analysis with *BEAST2.5.2 ^56^, specifying a strict clock model, HKY substitution model and Yule prior of the species tree. MCMC chains were run for 20 million generations of which the first 5 million generations were treated as burn-in and discarded. The resulting species tree placed *O. retusus* as being more closely related to *O. gibbosus* compared to *O. fuscus*, and the former species was therefore selected as closest outgroup in further analyses (Fig. S8). Maximum clade credibility trees for each individual window were obtained with TreeAnnotator. Trees were visualized with the *densiTree* function included in the *phangorn* v2.5.5 package in R v3.5.3. For all windows (including the *G* locus) we calculated the scaled maximum coalescent time for *O. gibbosus* by dividing the maximum node height of the *O. gibbosus* haplotypes by the height of the node that splits *O. gibbosus* from *O. retusus*.

### Structural variation between the morph specific alleles

Structural variations (SV) between the morph specific alleles were detected using PacBio data only ^57^ . PacBio reads originating from a pool of heterozygous hunched and homozygous flat offspring, all originating from the same family, were mapped separately to the heterozygote hunched genome assembly (see *Genome assembly* “Ogibo_pacbio_wtdbg2_het”) with ngmlr v.0.2.7 ^24^. We used Sniffles v 1.0.11 ^24^ to detect SV present uniquely in the heterozygote hunched or homozygote flat offspring pools. SV that were genotyped as heterozygote in the PacBio reads originating from the heterozygote hunched offspring and absent or homozygous in flat offspring were classified as morph specific and allowed us to assign them to either the hunched (G) or flat (g) allele. This SV analysis revealed that, except for a ∼25kb insertion, PacBio reads from the pool of homozygote flat individuals corresponded to the genomic region that we previously assigned as the *G*-locus (ctg337: 500kb – 800kb) (Fig. 2, Fig. S9), indicating that the *G*-locus of our reference heterozygote assembly represents the g-allele. In contrast, PacBio reads from the pool of heterozygote hunched individuals in contrast mapped only partially, with SV analysis revealing multiple insertions, including one of unresolved length (Fig. 2, Fig. S9). To further confirm that the most differentiated region in our heterozygote reference assembly corresponds to the g-allele, we aligned the homozygote flat genome assembly (see “*Genome assembly*”: Ogibo_pacbio_wtdbg2_flat) to our heterozygote reference genome assembly (see “*Genome assembly*”: “Ogibo_pacbio_wtdbg2_het”) with minimap2 ^39^. This alignment identified a single 760kb contig that fully aligned with the differentiated region on ctg337 and allowed us to accurately determine the structure of the flat allele (Fig. S10). Because we have no access to large numbers of homozygote hunched siblings for PacBio sequencing, we manually assembled 380kb of the G-allele based on G-allele associated reads extracted from the heterozygote hunched offspring pool. Hunched associated PacBio reads were visually identified based on (i) genotypes that segregated according to the expected genotypes of the resequenced individuals (see “*Whole genome resequencing, genotyping and sequence divergence*”), (ii) genotypes observed in resequenced homozygous hunched and flat individuals and (iii) SNPs/SV that were not observed in the PacBio reads originating from homozygote flat individuals (Fig. S8). PacBio reads were imported in Geneious v 11.0.5 (© Biomatters Ltd.) and mapped iteratively using the “map to reference” tool specifying a high sensitivity. Only reads with an overlap of at least 2kb were used to assemble the hunched allele.

Structural variations between the reconstructed hunched and flat allele were inspected using dotplots constructed by Geneious v 11.0.5 (© Biomatters Ltd.). Alignment of the assembled flat and hunched allele revealed multiple insertions in the hunched allele, often spanning several kb in size (Fig. S11). The sequence parts that were absent in the flat allele generally comprised non-unique sequences that mapped to multiple regions in the genome. Reads associated with the hunched allele were further clipped about 4kb upstream this ∼25kb flat deletion (ctg337: 643,370) at the point where our Sniffles v 1.0.11 ^24^ analysis detected an insertion of unresolved length (Fig. S9). Moreover, one insertion in the hunched allele could not be resolved because PacBio reads at both sides of the insertion did not overlap (Fig. S11).

Based on the presence of these potentially unresolved insertions in the hunched allele, we screened the entire genome for contigs showing signatures of being only present in hunched individuals. More precisely, we first tested this by screening the entire genome for contigs showing a strong signature of being uniquely present in hunched individuals. A contig was proposed to be uniquely present in hunched males if its coverage was significantly higher in hunched compared to flat individuals (*t* -test; P < 0.01) and if it contains positions that are consistently covered by short- and long reads from hunched individuals, but not in those from flat individuals (Fig. S12). To test for significant differences in contig coverage depths between the morphs, we first calculated for each resequenced individual (including the outgroup species *O. retusus* and *O. fuscus*) the coverage at both 2kb window and contig level with the pipeCoverage v1.1 tool (https://github.com/MrOlm/pipeCoverage). Coverages were subsequently scaled at individual level by dividing the coverage per window or contig by the median coverage of this individual across all windows. Differences in sequencing coverage between the eight resequenced hunched and eight flat individuals for each contig were then calculated as the log2FoldChange, being the log2 transformed average scaled coverage in the hunched individuals divided by the average scaled coverage in the flat individuals and tested by means of an unequal variance t-test. To test for the presence of contigs containing genomic positions consistently covered by hunched but not in flat, we first obtained sequencing depths at all genomic positions for each individual and the PacBio sequenced pools of hunched and flat offspring from the individual bam files with the ‘depth -a’ command in Samtools v0.1.20 ^50^. Positions having coverages of 0 in all resequenced flat individuals and the PacBio reads of the pool of flat offspring, but higher than 0 in all hunched individuals and PacBio reads of the pool of hunched offspring, were subsequently counted per contig using a custom awk script. Contigs matching both the criteria of lower coverage depth (*P*< 0.01) and missing positions were considered hunched-specific. To verify if this selection approach could result in false positives, we randomly shuffled morph phenotypes among individuals (100 replicates) and tested with the same procedure for the presence of positions showing a consistent absence in flat individuals. None of the reshuffling replicates recovered a single position matching this criterium and the probability that our selection includes false positives is < 0.01.

Lastly, we searched for a potential homologous flat allele by mapping the hunch-specific contigs to the remaining genome sequence with minimap2 ^39^ and retained mappings with a minimum mapping quality of 30. To test if the targets of these mappings point towards a potential homologous flat-specific sequence or to a repetitive region that is present in the genome of both morphs, we calculated the average coverage depths of the target regions with BEDTools v.19.1^58^ (bedtools coverage -mean -a “bedfile specifying the mapping targets” – b …).

### RNA sequencing

We extracted total RNA from five adult hunched males, five adult flat males, two subadult hunched males, two subadult flat males and six (four Gg and two gg genotypes) adult females (Table S1) using the RNeasy Plus Universal Mini kit (Qiagen). Before extraction, live specimens were frozen at -80°C and 2-3 legs were removed from the frozen specimens to genotype individuals at the G locus with the diagnostic marker (see “*PCR genotyping at the G-locus*”). Strand-specific Illumina sequencing libraries were prepared with the TruSeq Stranded Total RNA with Ribo-Zero HMR kit (Illumina), except for two adult *gibbosus* and *tuberosus* males for which the non-strand-specific TruSeq Total RNA with Ribo-Zero kit (Illumina) was used. Libraries were sequenced on an Illumina NovaSeq platform (Macrogen) resulting in between 20M and 29M paired-end reads of 150bp per individual (Table S1).

### Repeat content

We screened the assembly (“Ogibo_pacbio_wtdbg2_het”) for repetitive elements with RepeatMasker v1.295 ^59^ specifying ‘Arachnida’ as species and hits to transposable elements (TEs) were assembled as complete elements with OneCodeToFindThemAll ^60^. A library of *de novo* repetitive elements was compiled with RepeatScout v1.0.5 ^25^ and used, together with the library of known repeats, to soft-mask repetitive sequences in the genome assembly. The distribution of repeats across the genome was estimated by calculating the number of TEs or the proportion of repetitive sequences in 3Mb windows with a customized Python script using the General Feature Format (GFF) files of the assembled TE’s (OneCodeToFindThemAll) or the total repeat content (Repeatmasker) as input files.

### Structural and functional gene annotation

*Ab initio* gene prediction was performed with the Braker2 pipeline ^61–64^ using the RNA sequencing reads as extrinsic information. Briefly, we first mapped RNA sequencing reads (stranded only) with HiSat2 v2.1.0 ^65^ to the “Ogibo_pacbio_wtdbg2_het” genome assembly. Resulting bam files were then used as input to train GeneMark-ET ^66^ with the spliced alignment information contained within the aligned RNA reads. Predicted genes are then used as input to train AUGUSTUS ^67^, resulting a final set of predicted gene models. Functional annotation of the predicted gene models was performed through similarity searches of the translated gene sequences using both BLASTP ^68^ against the UniProtKB/SwissProt ^69^ protein database using an E-value cutoff < 0.0001 (downloaded 4/3/2020) and InterProScan v. 5.28-67 ^70^. We further generated a *de novo* transcriptome assembly with Stringtie v2.0.3 ^71^ using the aligned RNAseq data, which was used for the manual curation of genes of interest.

For all predicted genes located on the hunch-specific sequence, unique presence in hunched males was confirmed by testing if coverage depths of the exon sequences, obtained with BEDTools v.19.1^58^ (bedtools coverage -mean -a “bedfile specifying the gene introns” – b …), was significantly higher in hunched versus flat males (unequal variance t-test, *P* < 0.01).

We searched the entire genome for genes paralogous to those located on the hunch-specific sequence by mapping their predicted coding sequences sequence to those of all genes located on the remaining genome sequence with Exonerate v2.4.0 ^72^.

All functionally annotated genes on the hunch-specific sequence were manually curated by investigating the presence of premature stop-codons and partial lengths by comparing the predicted gene models with aligned RNAseq data (if exptressed) and Exonerate v2.4.0 ^72^ aligned CDS of their closest paralog located outside the hunch-specific sequence and CDS of most similar reference protein sequences in the UniProtKB/SwissProt ^69^ database by means of the GenomeView (https://genomeview.org/) genome browser.

### Differential gene expression (DE) analysis

We obtained raw individual RNA sequencing read counts for each predicted gene with *Stringtie v2*.*0*.*3* ^71^ and calculated normalized read counts and gene expression differences with the Deseq2 v1.30.0 ^73^ package in R v3.5.2 (R development Core Team). Differences in gene expression between the male morphs were obtained by estimating the log_2_-fold changes in transcript abundance, and their associated P-values (Benjamini-Hochberg adjusted for multiple testing), between the adult hunched and adult flat males. The same procedure was used to obtain sex-specific differences in gene expression and was calculated for each male morph type separately by comparing both (i) hunched (Gg) males versus Gg females or (ii) flat (gg) male versus all females.

We tested if the expression of genes located on the hunch-specific sequence (n = 195 predicted genes) are significantly more male-biased by comparing their average as well as median sex-specific log_2_FC with those obtained in 10,000 random samples of 195 genes located in the remaining genome sequence by means of a custom R script. A similar procedure was used to test if the expression of the closest paralogs (n = 149) of the genes located on the hunch-specific sequence are significantly more male-biased compared to those located in the remaining genome sequence.

### DMRT and AChE analysis

We identified *Doublesex/mab3 related transcription factor* (*Dmrt*) related genes in the *O. gibbosus* genome by two complementary methods. We first screened the entire genome for Dmrt related genes by tBLASTn searches using the Dsx1, Dsx2, Dmrt11E, Dmrt93B, Dmrt99B and three Dmrt_like sequences of the spider *Parasteatoda tepidariorum* ^20,74^ and the Dsx1, Dmrt11E, Dmrt93B and Dmrt99B sequences of *Drosophila melanogaster* as query sequences and the genome as database. Significant hits (E-value < 3E-07; Table S4), all involving the DM DNA binding domain, were found at 12 distinct genomic locations. Second, we searched our InterProScan annotation results of the Braker2 predicted genes (see “Structural and functional gene annotation”) for genes containing a DM DNA binding domain, which yielded 11 genes. All 12 regions obtained from our tBLASTn searches were included in the 11 predicted Dmrt genes of InterProScan, with one predicted gene (OgDmrt_g20232) containing two DM domains. A final manual curation of all Dmrt genes, partially based on transcriptome data obtained from RNAseq, allowed us to reconstruct the protein sequences of the 11 *Dmrt* paralogs. All contained the M DNA binding domain (conserved domain search with CDD v3.17 ^75^; E < 3.6E-19).

*Dmrt* paralogs were placed into the pan-arthropod phylogenetic framework developed by Panara *et al*. ^74^ by comparing their phylogenetic relationship with those of the spider *Parasteatoda tepidariorum*. To assess the relationships of the two DM domains contained within OgDmrt_g20232, the coding sequence was split at a 5.7kb intron separating the two domains. Predicted sequences of all *O. gibbosus*, as well as the *P. tepidariorum* paralogs, were aligned with COBALT ^76^ and phylogenetic relationships were estimated based on the 195 aligned residues (including 47 residues in the DM domain). Maximum likelihood phylogenetic analysis was conducted with RAxML v8.2.11 ^77^ specifying a fixed WAG substitution model and gamma distributed rate variation across sites.

We reconstructed the evolutionary history of the three *Acetylcholinesterase*-like genes in the *gibbosus* sequence by comparing their sequence with those of all putative *Acetylcholinesterase*-like genes present in the genome of *O. gibbosus*, as well as those present in the genome of the spider *Parasteatoda tepidariorum* 20. Genes were selected by performing BLASTp searches of the three *Acetylcholinesterase*-like genes in the hunch-specific sequence against the predicted gene sequences of *O. gibbosus* (E-value < 1E-34) and the official gene set of *P. tepidariorum* (E-value < 1E-12). Phylogenetic relationships among the amino acid sequences were reconstructed using the same alignment and phylogenetic analysis procedure as for *Dmrt*.

## SUPPLEMENTARY FIGURES

**Fig S1.**
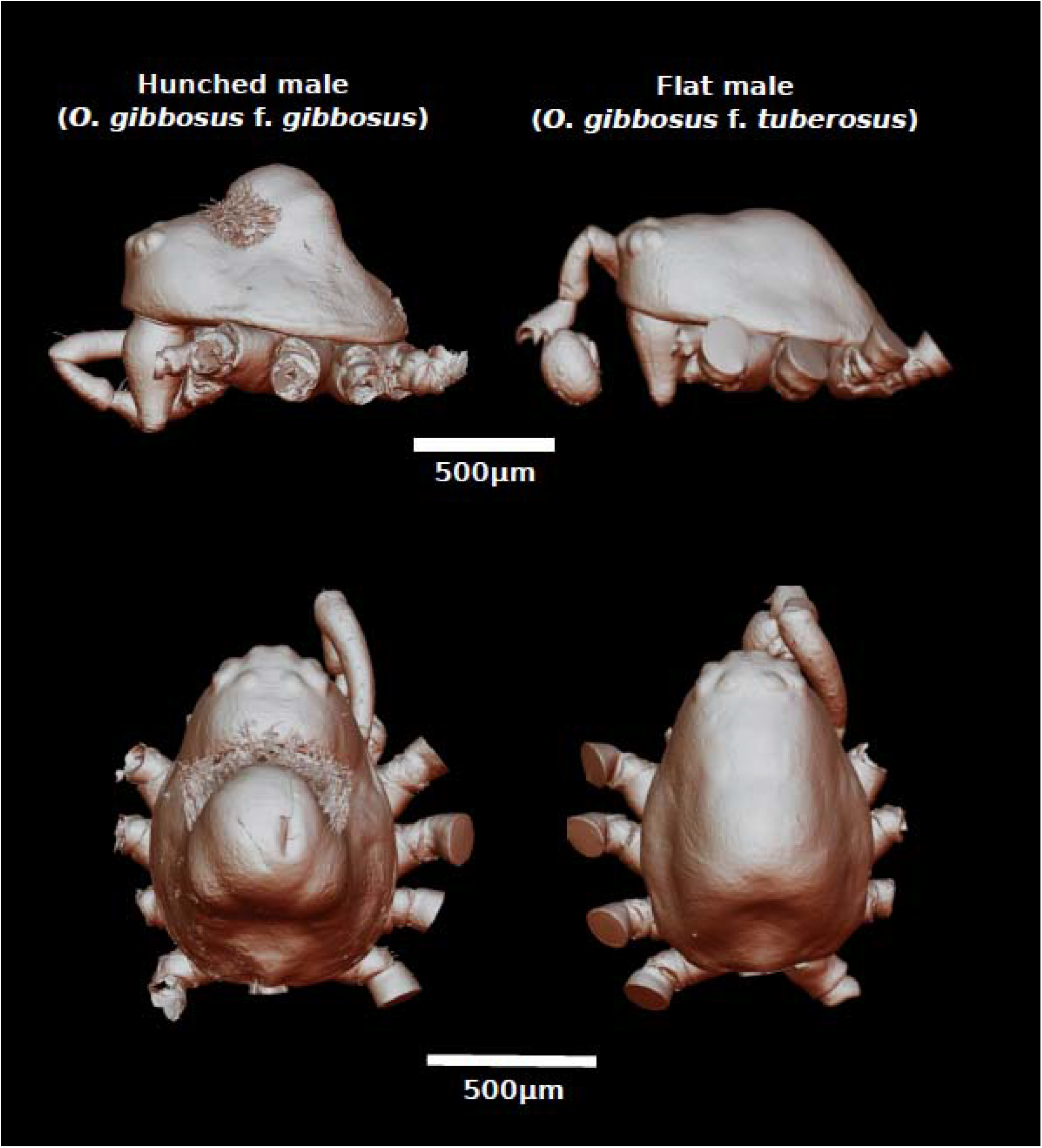
High-resolution micro-CT scanning of the carapaces of the hunched and flat male morph of *Oedothorax gibbosus*. Legs and abdomen were removed to visualize the difference in cephalic structures.

**Fig S2.**
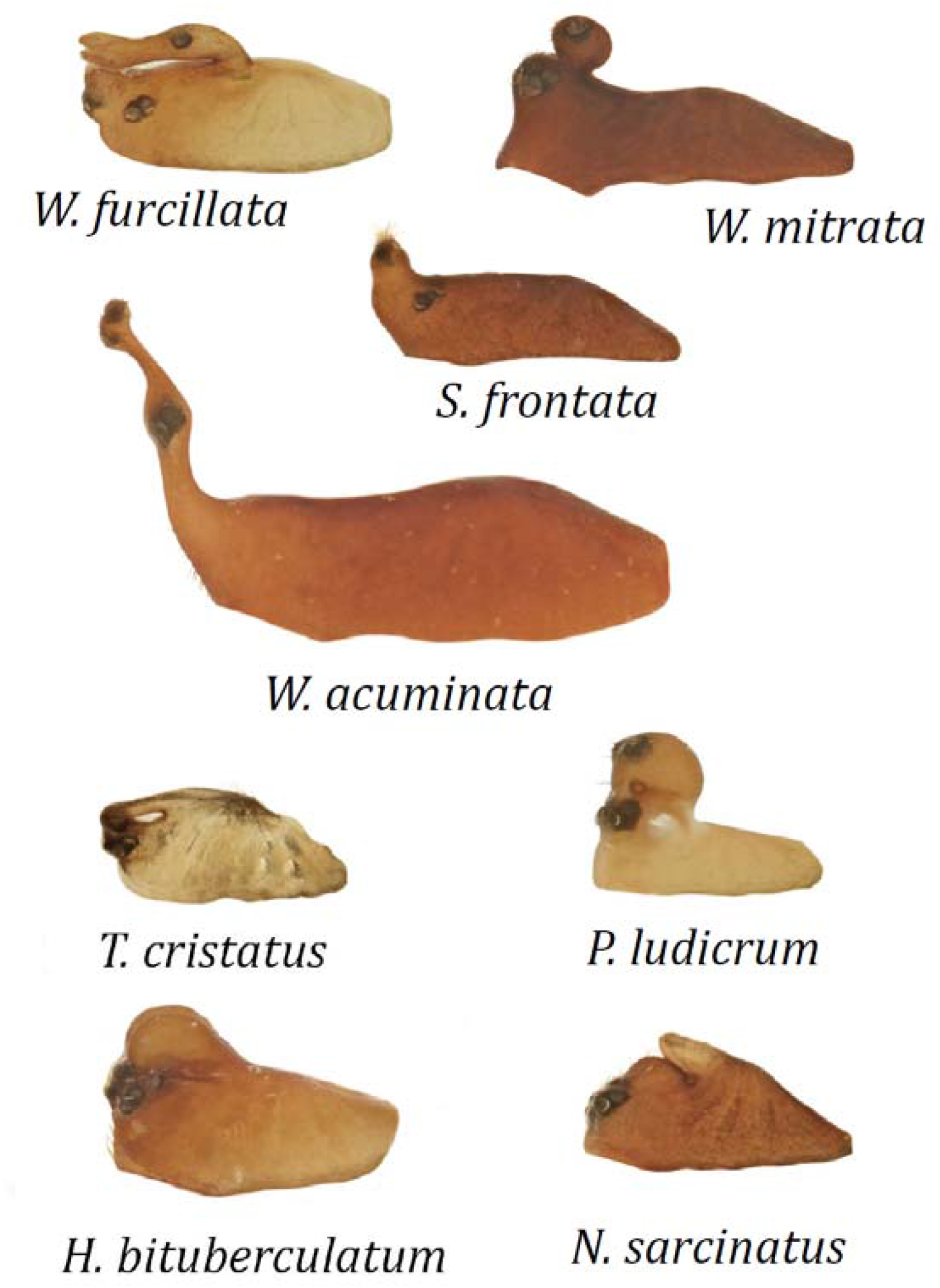
Examples of male carapaces within the subfamily of dwarf spiders (Erigoninae). Depicted species are *Walckenaeria furcillata, Walckenaeria mitrata, Savignya frontata, Walckenaeria acuminata, Trematocephalus cristatus, Peponocranium ludicrum, Hypomma bituberculatum and Notioscopus sarcinatus*.

**Fig S3.**
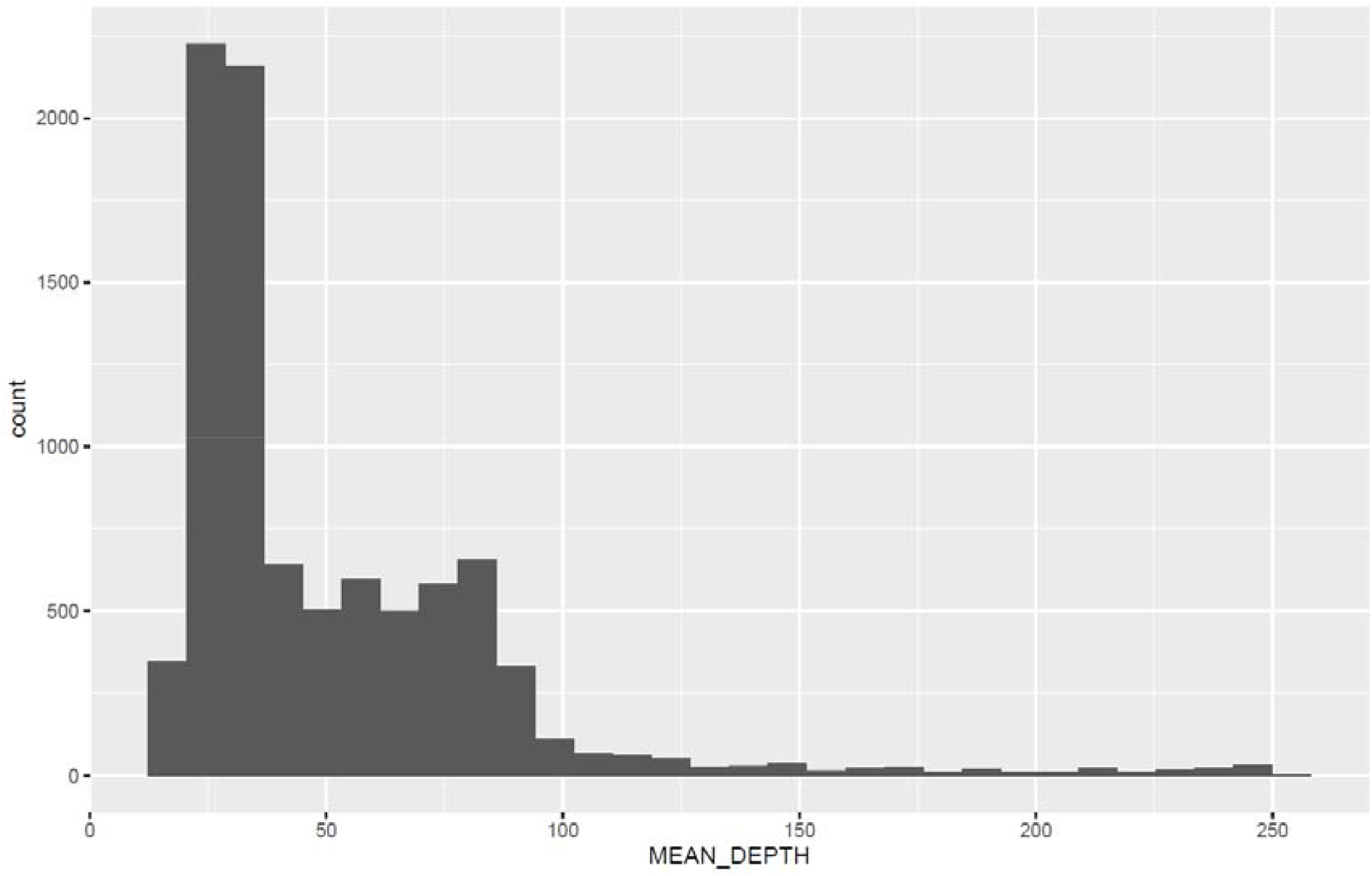
Distribution of the mean depth of the 9179 filtered SNP markers obtained by RADtag sequencing of offspring from a cross between a hunched male (W834) and a female (W828). SNPs with depths > 110 were excluded for linkage map construction.

**Fig S4.**
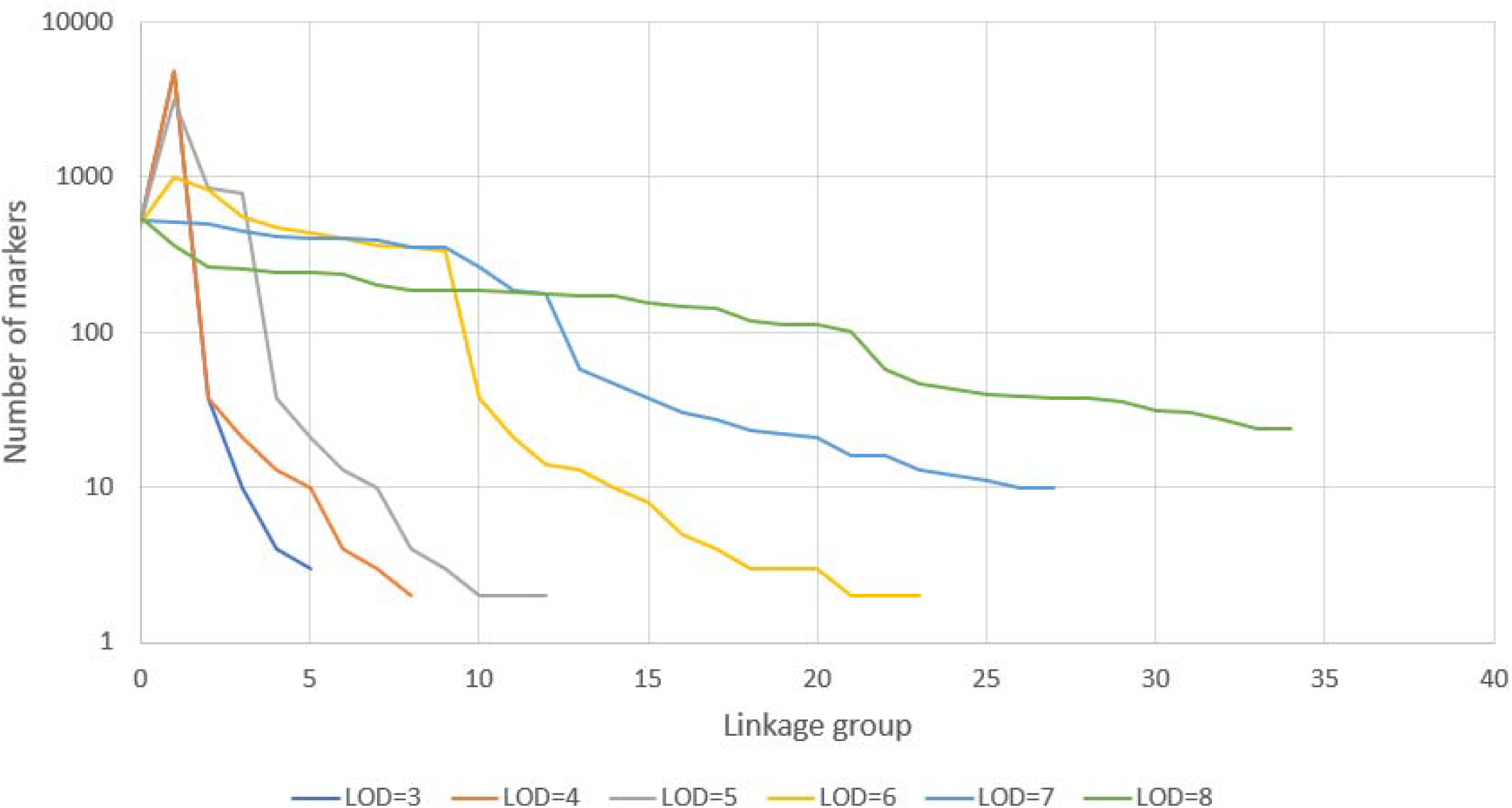
Distribution of the number of markers per linkage group depending on the defined two-point LOD score to group markers into linkage groups.

**Fig S5.**
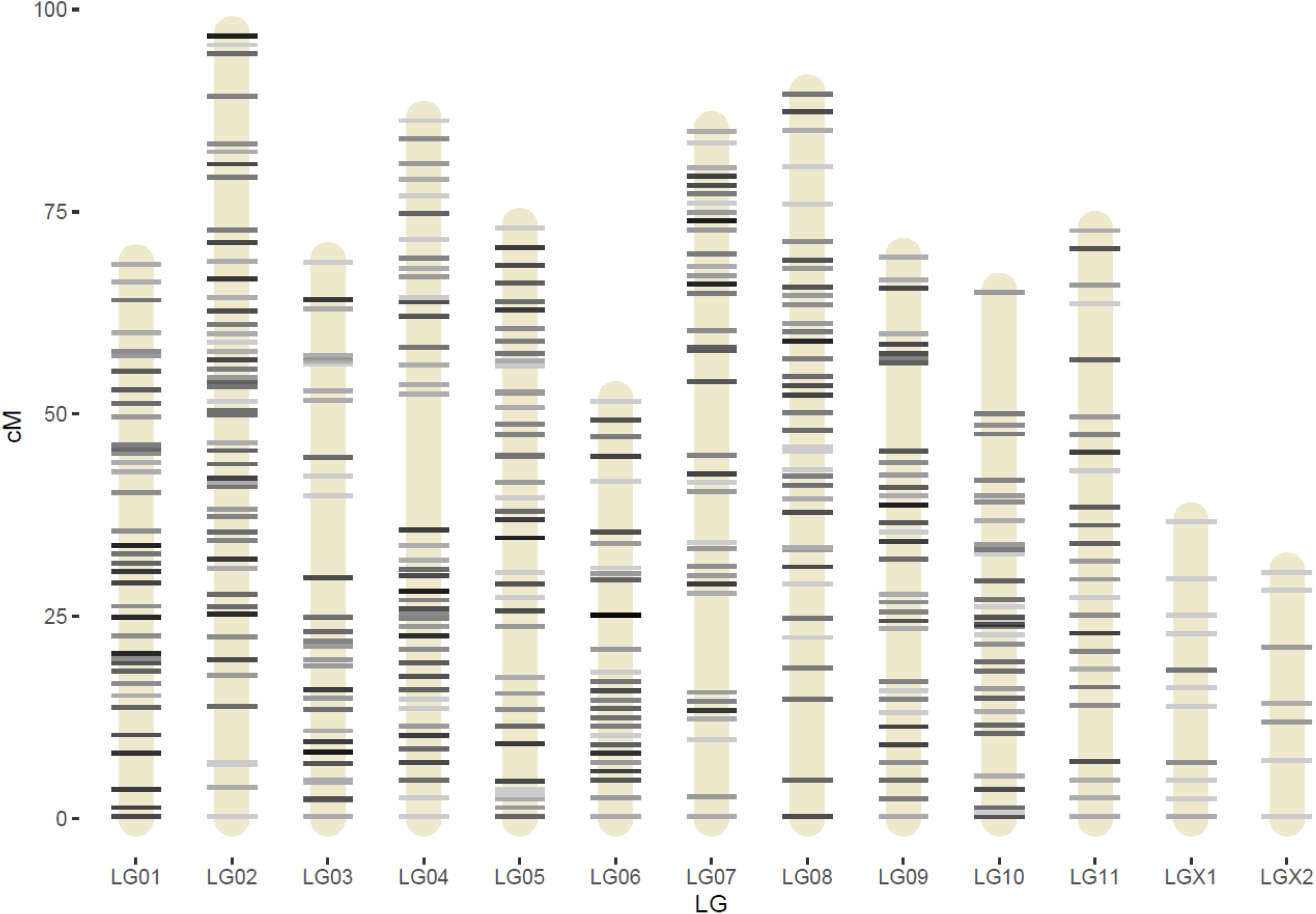
Linkage map of *O. gibbosus*.

**Fig S6.**
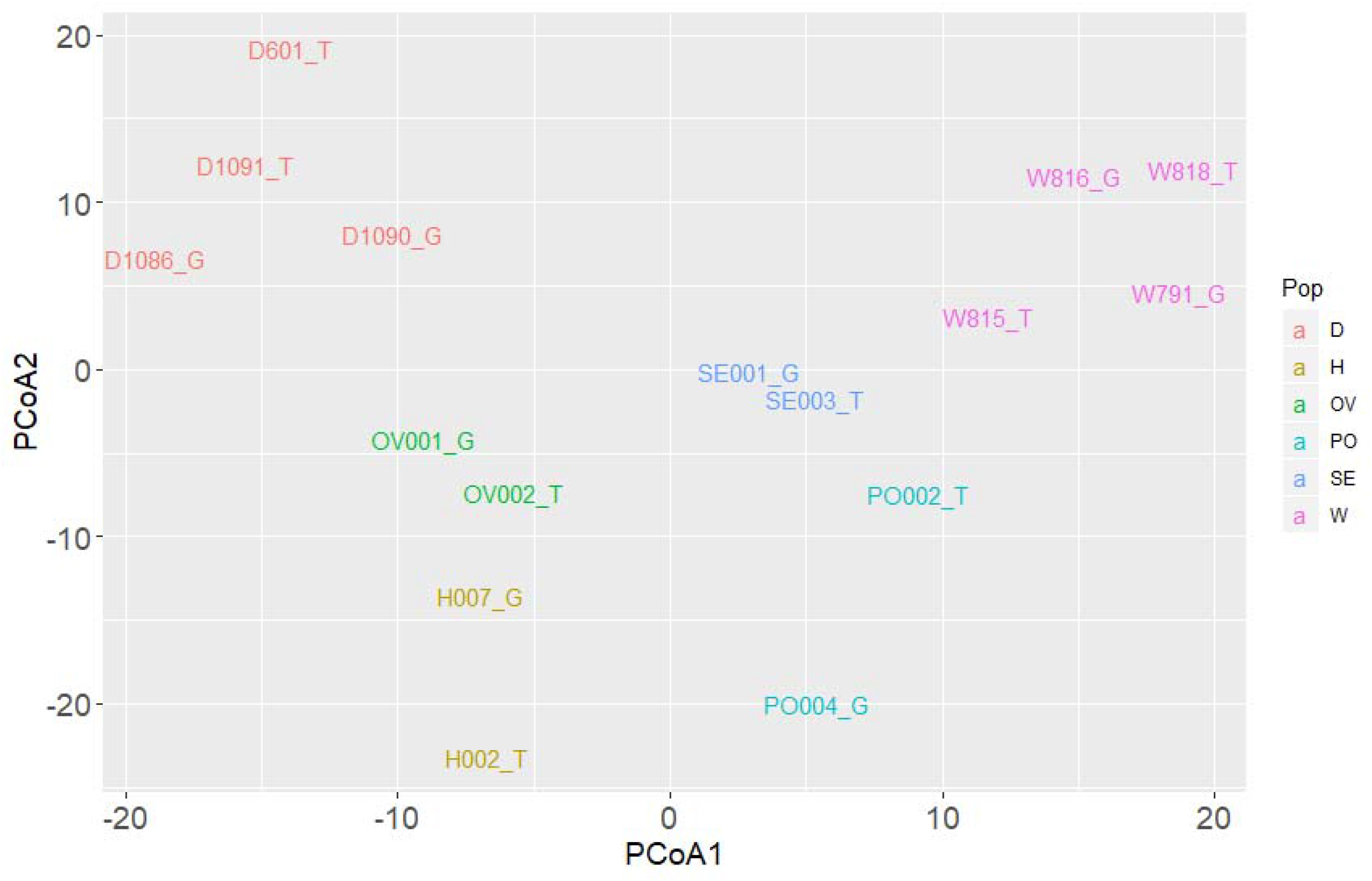
Principal coordinate analysis (PCoA) of the resequenced *O. gibbosus* individuals based on SNPs with high quality genotypes (GQ > 30) in all 16 individuals and thinned to 1SNP per 20kb (39,366 SNPs in total). Hunched and flat males are labelled with “_G” and “_T” respectively.

**Fig S7.**
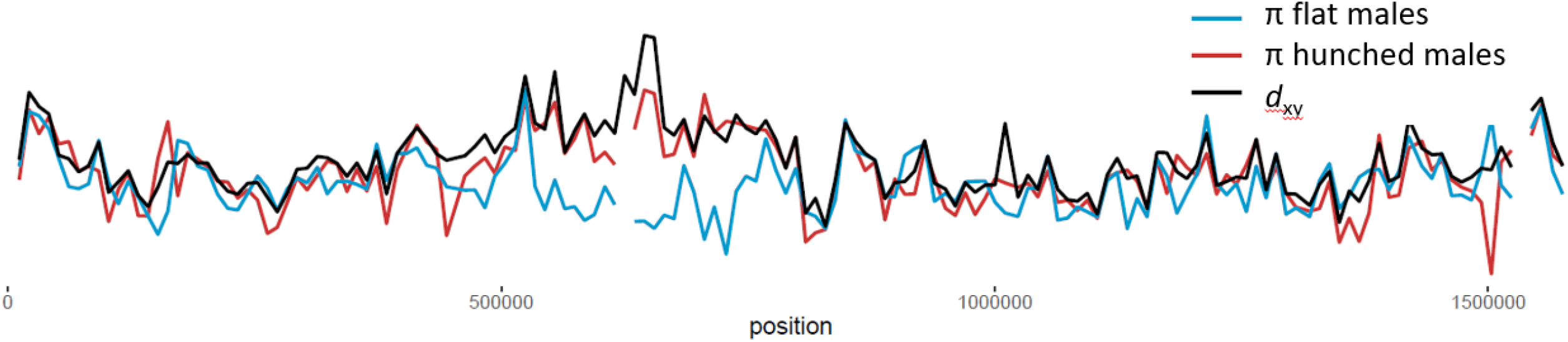
Patterns of nucleotide diversity (π) of the resequenced flat (blue) and hunched (red) individuals and sequence divergence (*d*_*xy*_) between the two morphs along 10kb windows at scaffold “ctg337”.

**Fig S8.**
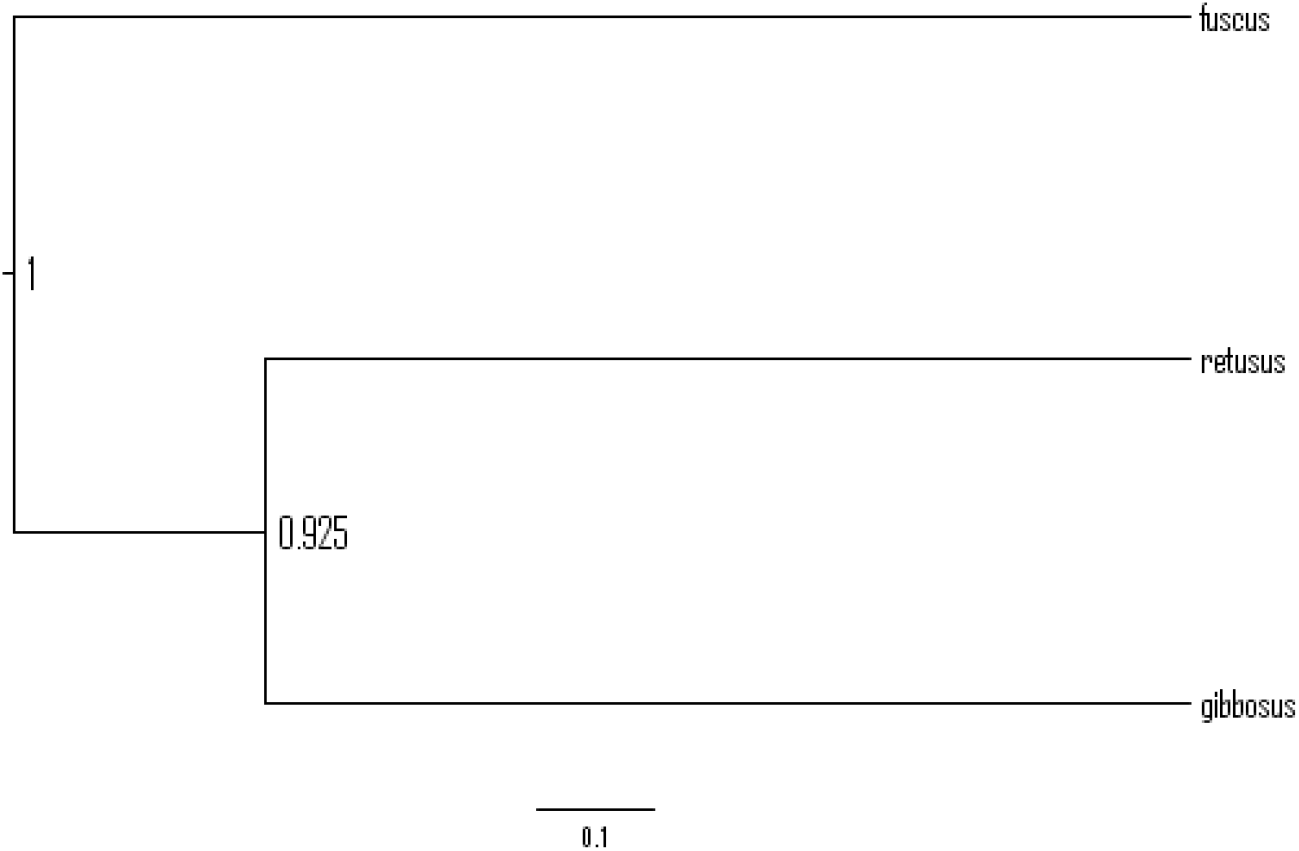
Species tree depicting the relationship between Oedothorax gibbosus and *O. retusus* and *O. fuscus* as inferred with *BEAST2 based on a set of 150 randomly selected 20kb windows.

**Fig S9.**
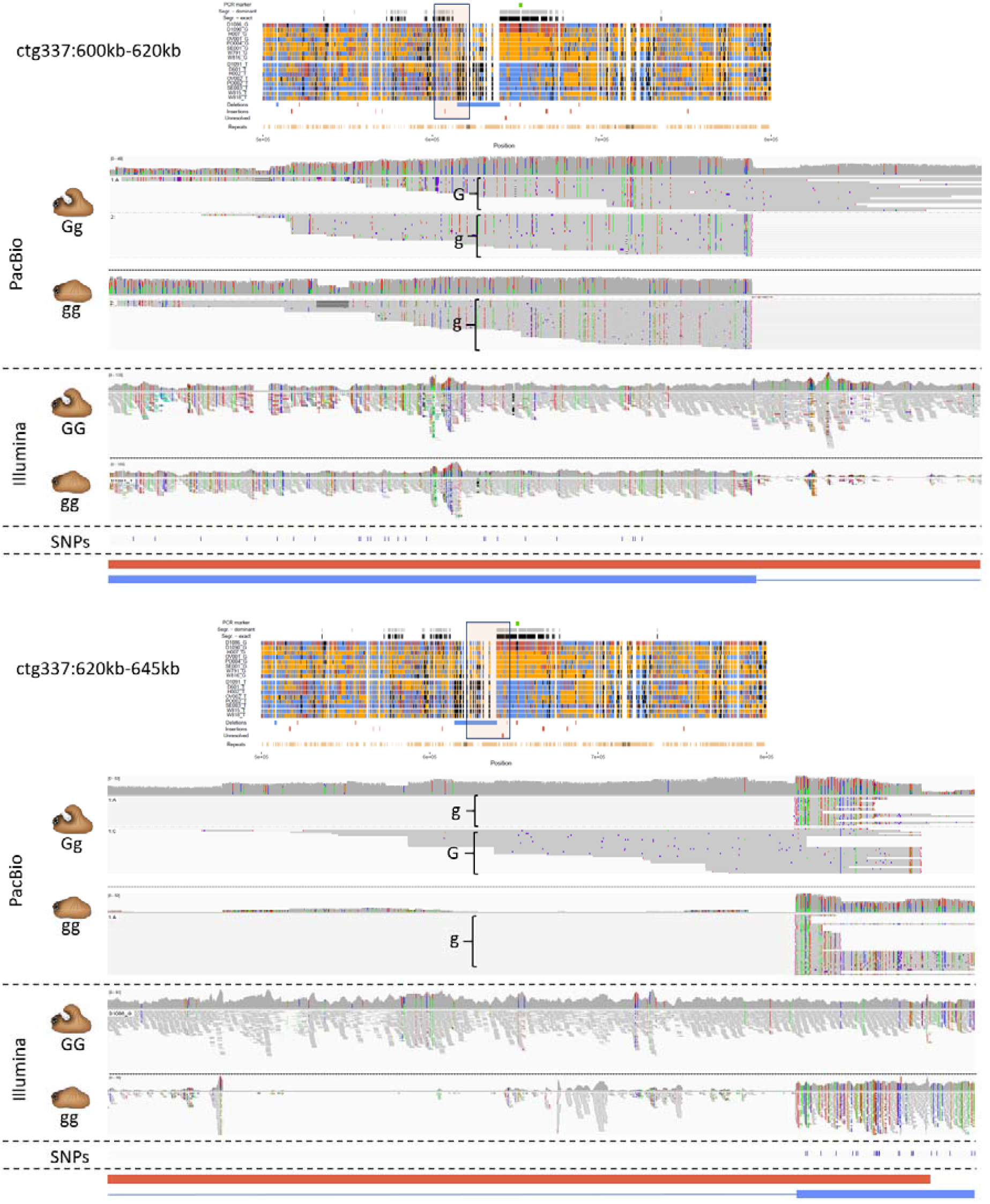
IGV screenprints of mapped PacBio long-reads (heterozygote hunched [Gg] and homozygote flat [gg]) and Illumina short-reads (homozygote hunched [GG] and homozygote flat [gg]), mapped around the ∼25kb deletion in the flat (g) allele (Ogibo_pacbio_wtdbg2_het: ctg337:600kb – 645kb). Red and blue bars below depict proposed structure of the hunched (G) and flat (g) allele in this region respectively. Upper and lower panel depict the sequence part upstream and downstream the ∼25kb indel.

**Fig S10.**
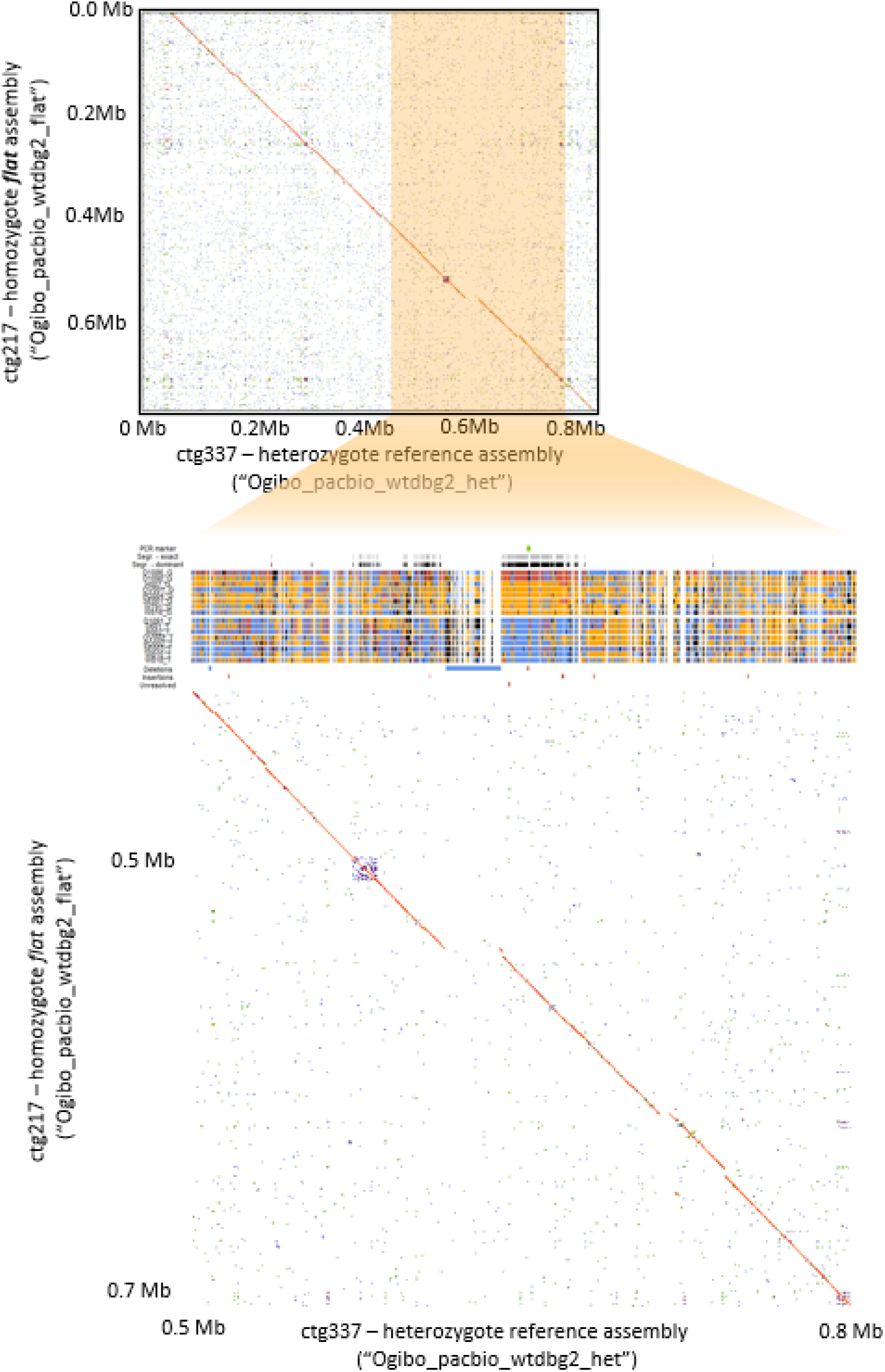
Minimap2 alignment of the heterozygote hunched assembly (“Ogibo_pacbio_wtdbg2_het”) with the homozygote flat assembly (“Ogibo_pacbio_wtdbg2_flat”) identifies a contig (Ogibo_pacbio_wtdbg2_flat: ctg217) that aligns across its entire length with the morph determining locus on ctg337.

**Fig S11.**
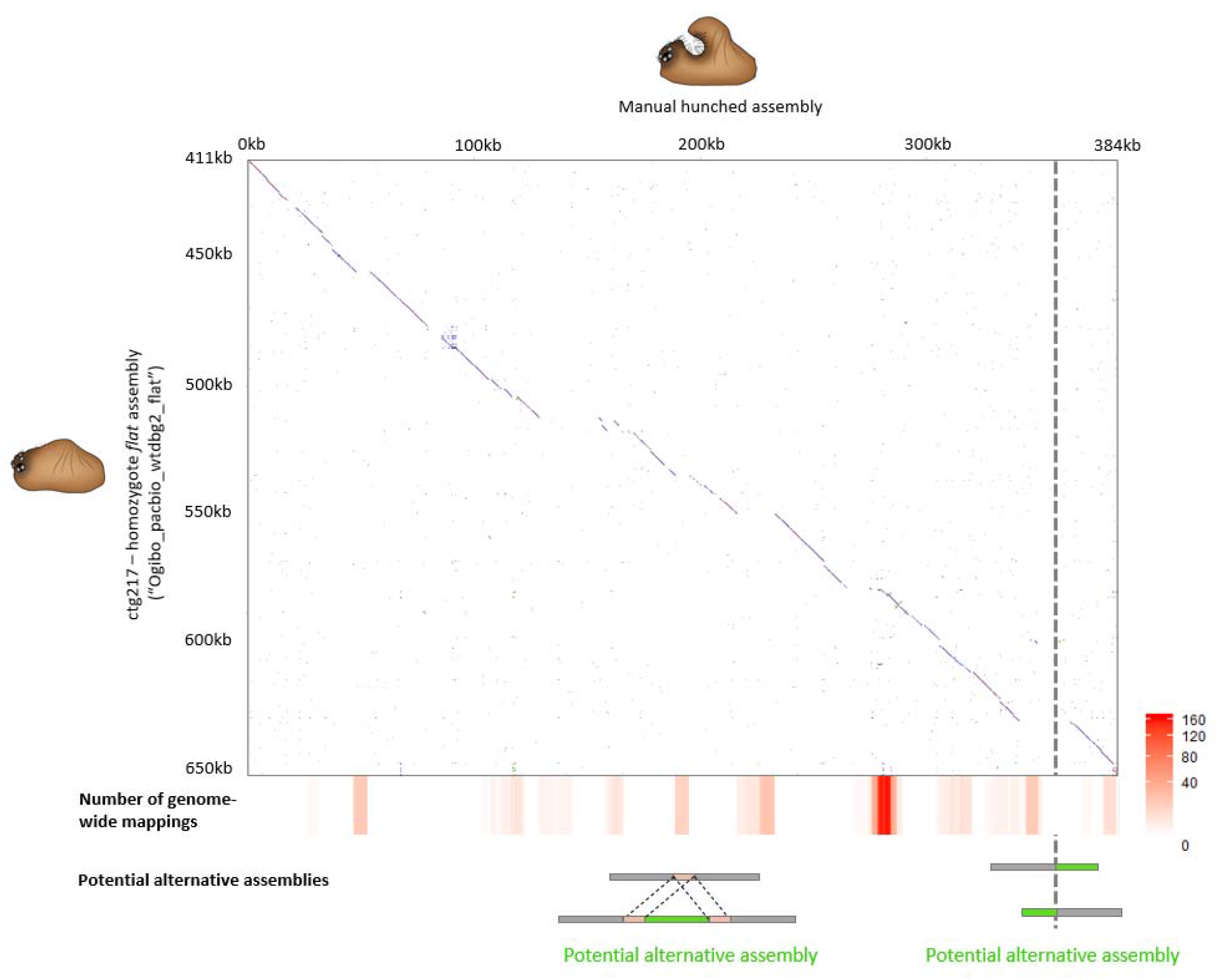
Minimap2 alignment between the flat-determining allele (Ogibo_pacbio_wtdbg2_flat: ctg217) and the manually assembled hunched-determining allele at the morph determining locus (ctg337: 500kb – 800kb in the heterozygote reference assembly). Dotted vertical line indicates an unresolved part in the hunched-determining allele due to the presence of non-overlapping reads and was artificially concatenated. Red colored bars below alignment graph depicts the repetitiveness of the sequence (>5kb) throughout the genome, expressed as number of times the sequence mapped to other regions in the genome. Bottom row illustrates how repetitive regions (centre) and non-overlapping reads (right) may result in ambiguous assemblies in the hunched allele, potentially leaving hunched-specific sequences (green) undetected.

**Fig S12.**
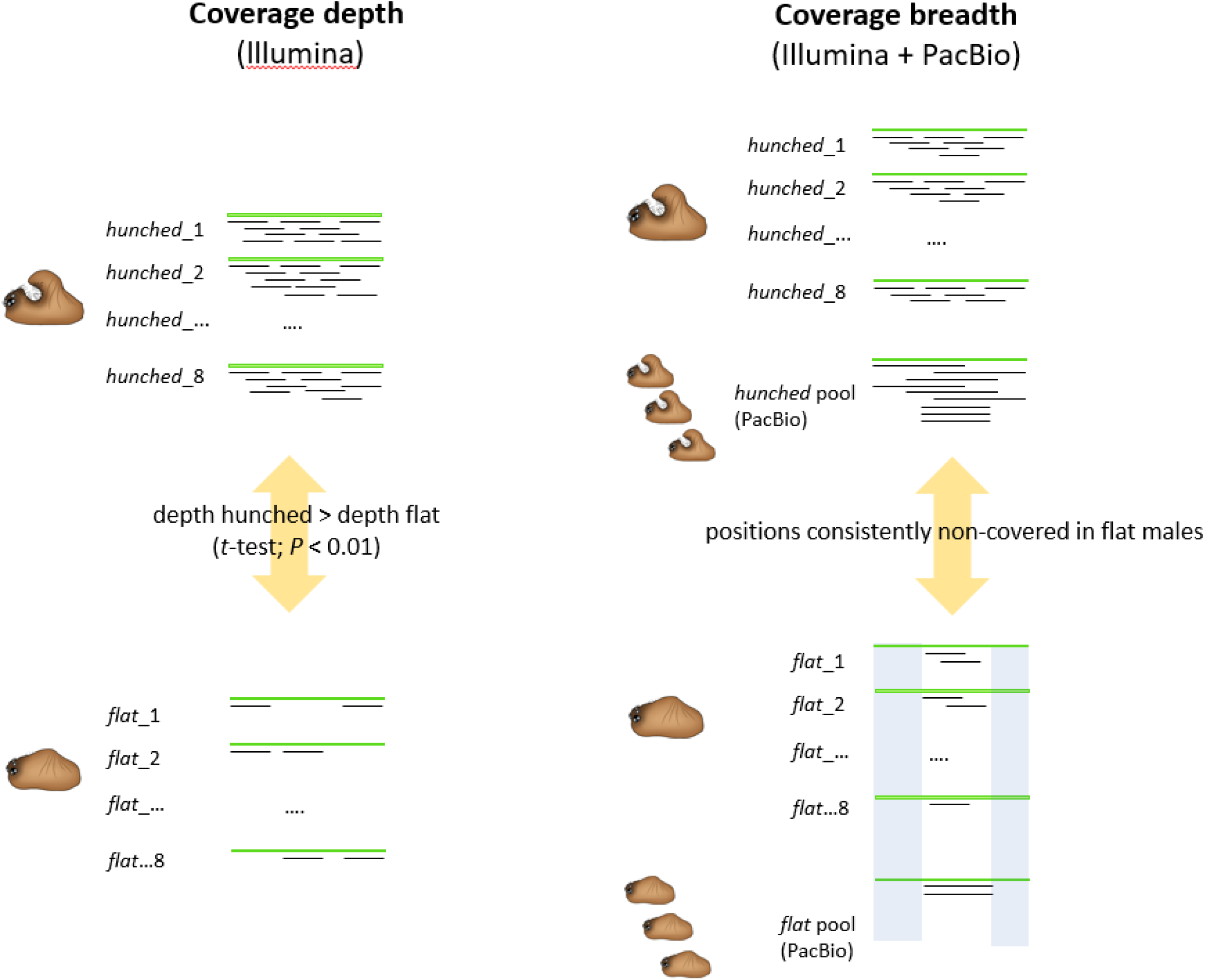
Strategy to identify contigs that are only present in the hunch-determining allele. Contigs having (i) significant lower coverage depths in the eight resequenced flat individuals compared to the eight resequenced hunched individuals (left panel) and (ii) positions that were only covered by Illumina short-reads in all resequenced hunched individuals and the PacBio reads of the pool of hunched offspring, but in none of the flat individuals and the pool of flat offspring (Right panel, indicated in light blue) were considered hunch-specific.

**Fig S13.**
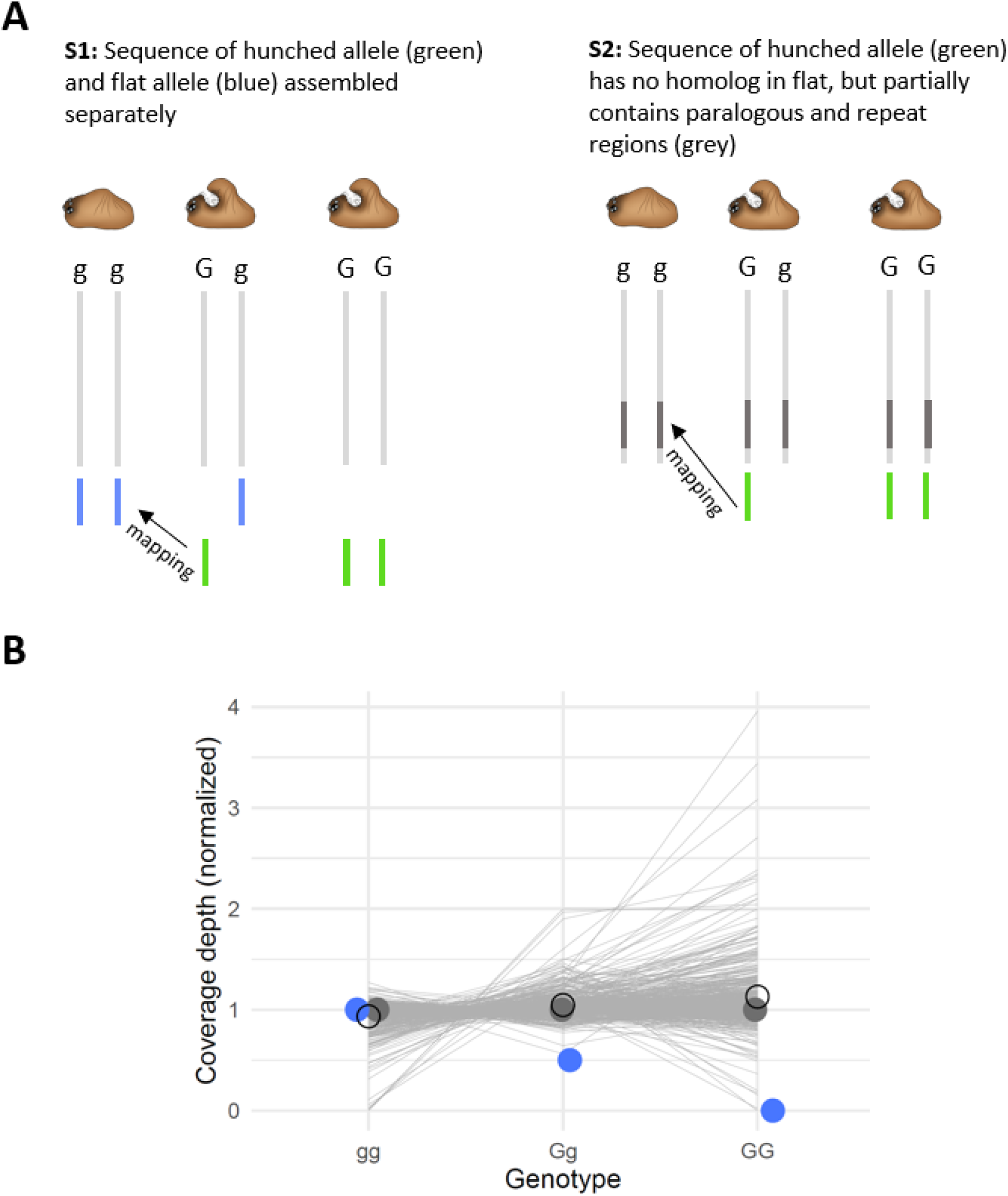
(A) Mapping of the contigs only present in the hunched allele to the remaining genome sequence could either represent (i) mappings to a potential homologous flat-determining allele (blue) assembled as a separate set of contigs (S1) or (ii) mappings to paralogous or repeated regions outside the *G*-locus (dark grey) (S2). (B) Coverage depths of the mappings presented in panel A for homozygote hunched (‘GG’; N = 2), heterozygote hunched (‘Gg’; N = 6) and homozygote flat (‘gg’; N = 8) individuals. Open circles show the average observed coverages depths. Blue and dark grey dots show the expected coverage depths of the mapping targets under scenario S1 and S2 in panel A respectively.

**Fig S14.**
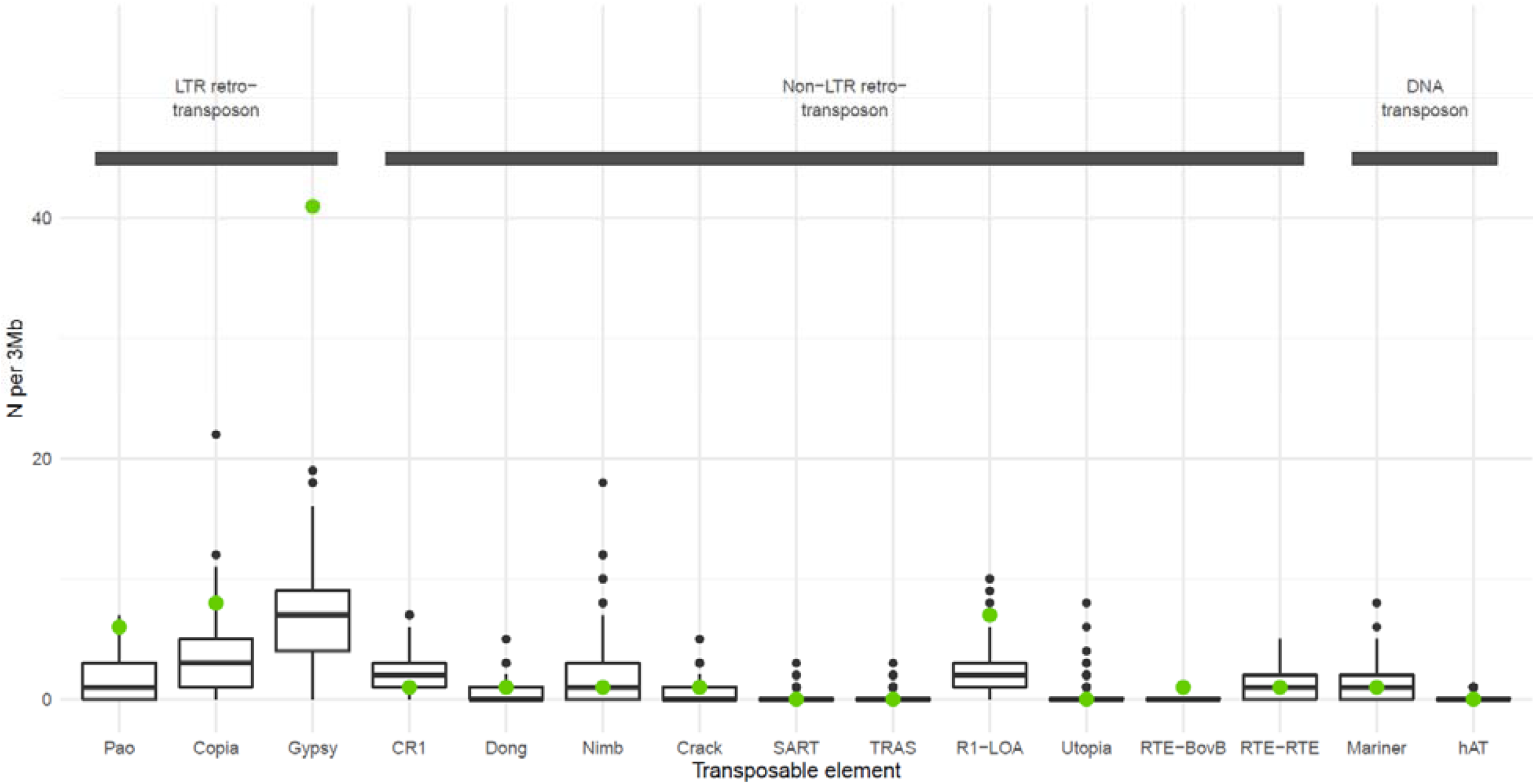
Distribution of the number of the 15 identified transposable elements (TE with names given below) in 3Mb windows across the genome. Green dots show the observed number of TE’s on the hunch-specific sequence.

**Fig S15.**
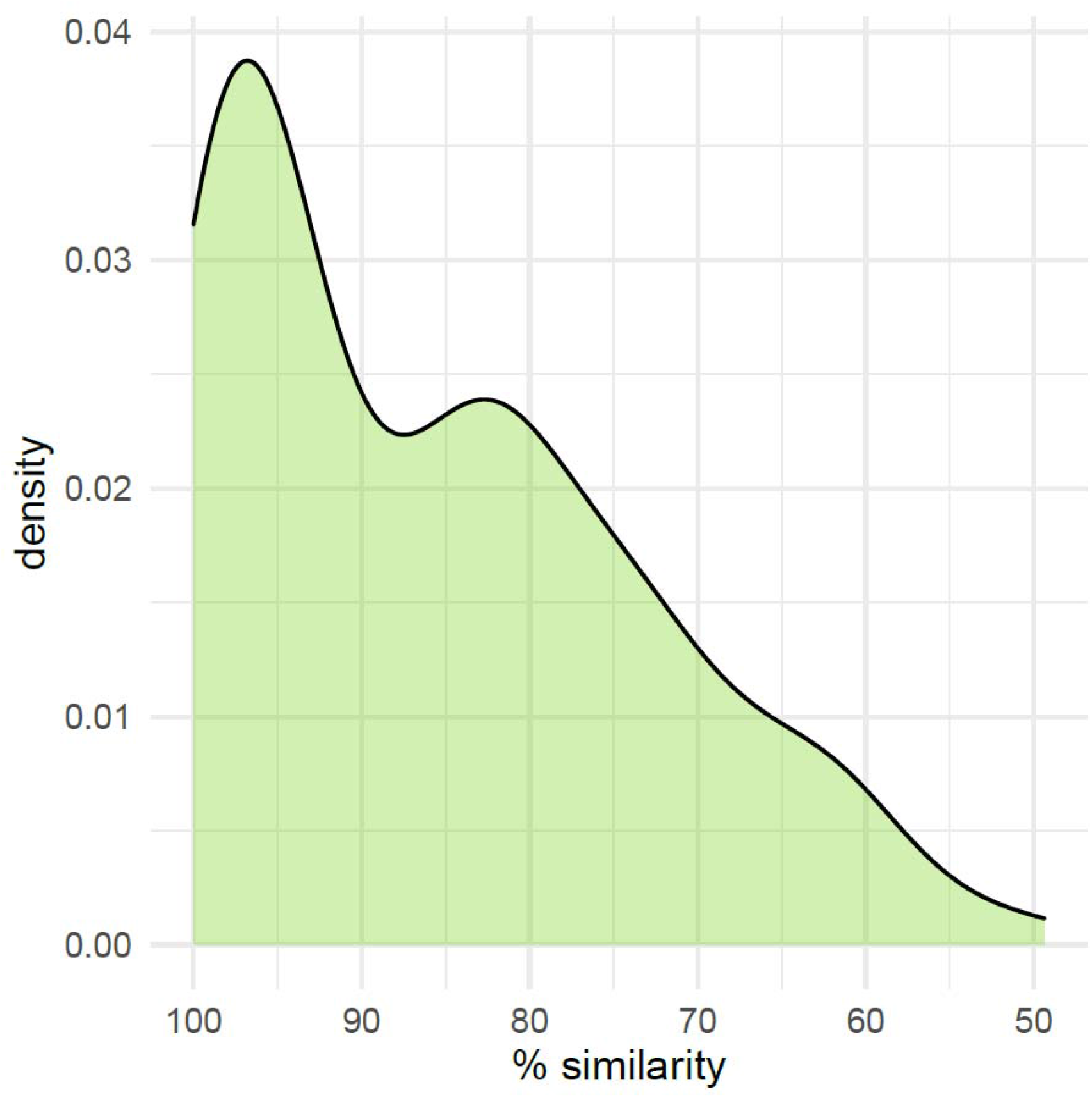
Density distribution of the % similarity of the encoded protein sequences on the hunch-specific sequence with those of their closest paralogs located outside the hunch-specific sequence (N = 149).

**Fig S16.**
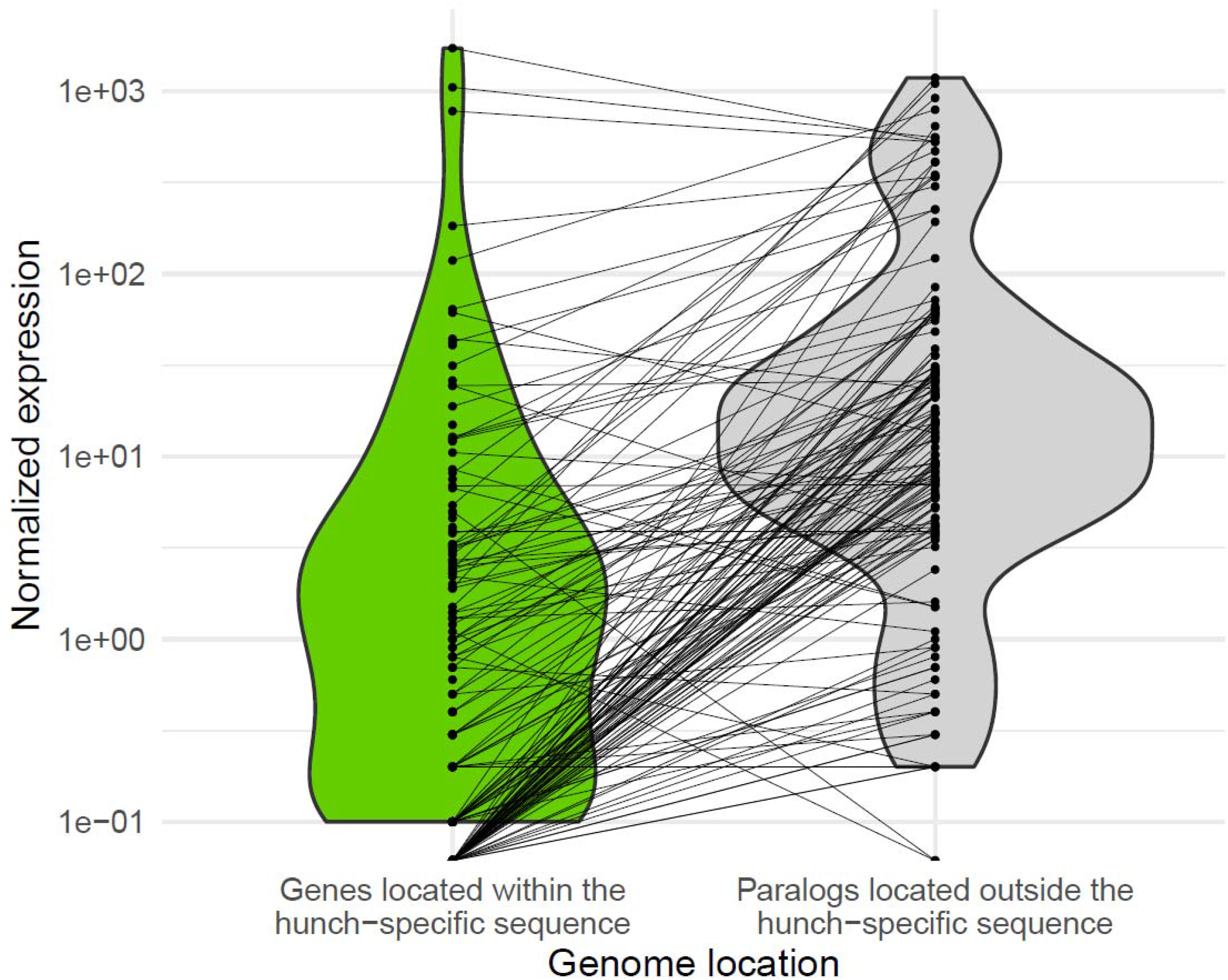
Comparison of the distribution of normalized gene expression of coding sequences (CDS) located on the hunch-specific sequence and their closest paralogs located outside this sequence fragment. Dots represent individual CDS connected with their closest paralog.

**Fig S17.**
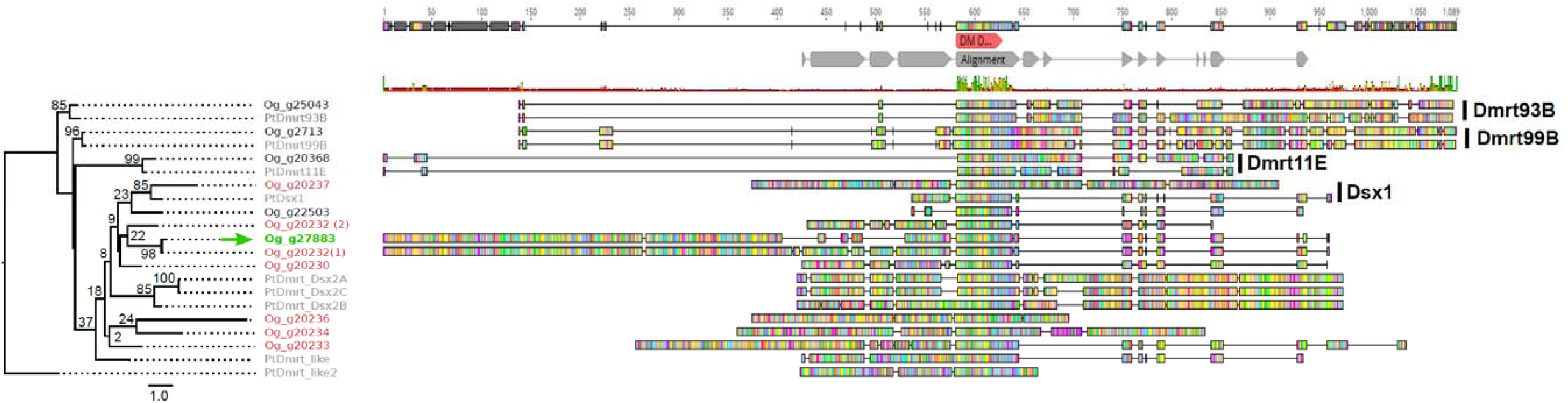
Alignment (COBALT) and maximum likelihood tree of the encoded protein sequences of all eleven Dmrt genes of *O. gibbosus* (Og) and those of the spider *Parasteatoda tepidariorum* (*Pt*). The tree was constructed based on the COBALT alignment (grey bars above sequences), with distances computed with RAxML v. 8.2.11 using the WAG substitution model and gamma distributed rate variation among sites. Red bar above alignments shows the location of the DM domain. Bootstrap values are presented next to the branches. The *Dmrt* gene located on the hunched fragment is indicated in green. *Dmrt* genes clustered on ctg151 are indicated in red.

**Fig S18.**
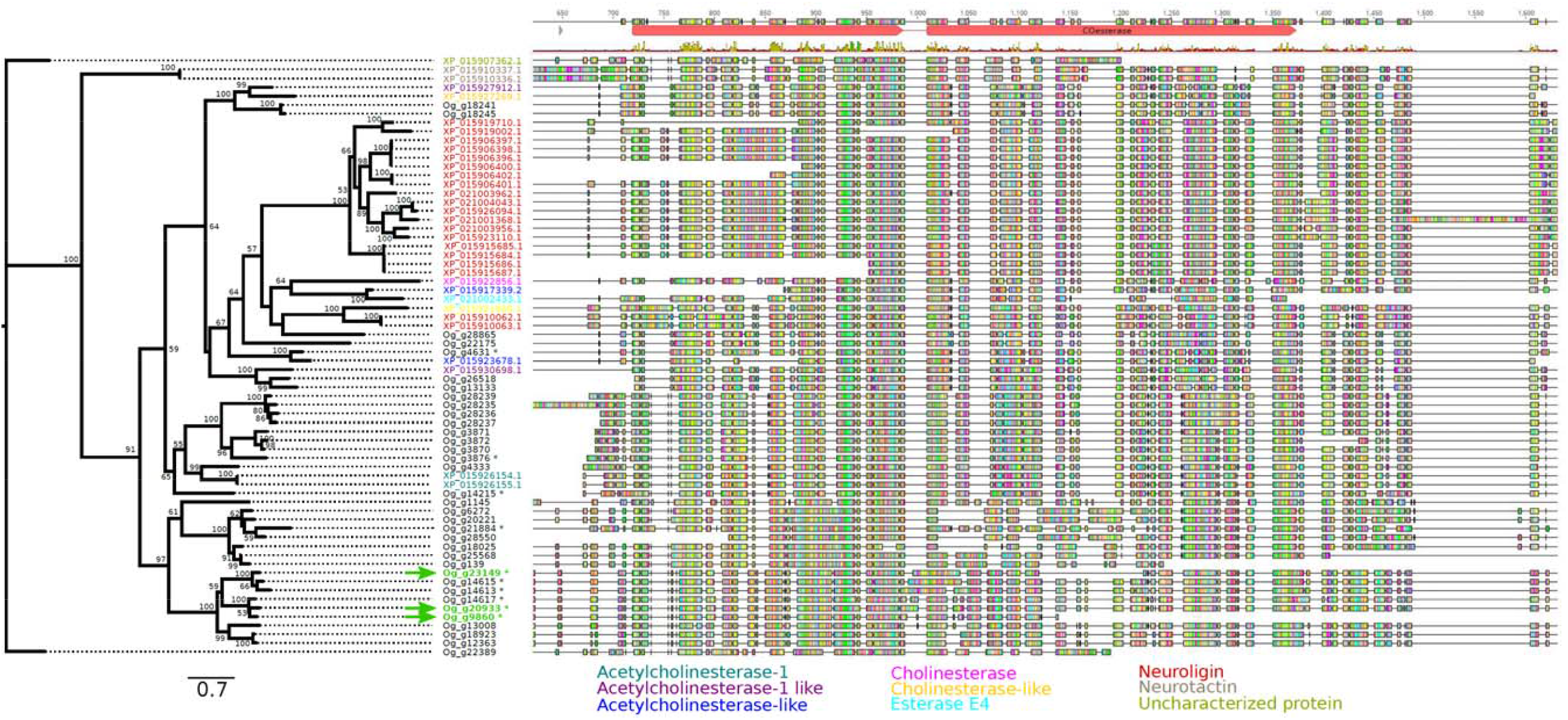
Alignment (COBALT) and maximum likelihood tree of the encoded protein sequences of all *Acetylcholinesterase* (*AChE*) related genes of *O. gibbosus* (“Og_x”) and those of the spider *Parasteatoda tepidariorum* (“XP_x”). The tree was constructed on the COBALT alignment with RAxML v. 8.2.11 using the WAG substitution model and gamma distributed rate variation among sites. Bootstrap values are presented next to the branches. The AChE genes located on the hunch-specific sequence are indicated with a green arrow, while those located outside the hunch-specific sequence are indicated in black. Sequence names of P.tepidariorum are color-coded according to their predicted function.

**Fig S19.**
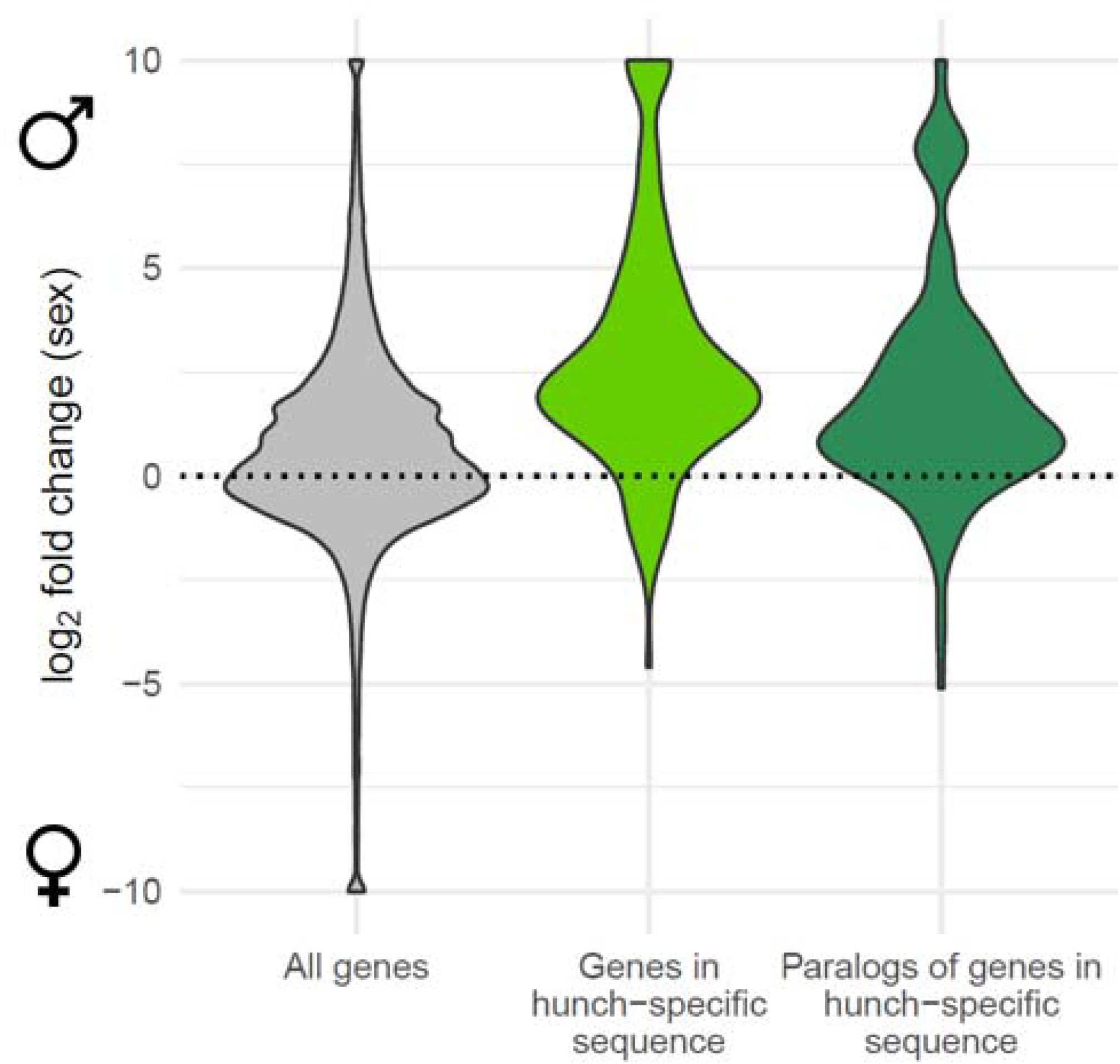
Distribution of the log_2_ fold change in gene expression between Gg males (hunched males, n = 5) and Gg females (n = 4) across all predicted genes (left, grey violin), genes located within the hunch-specific sequence (central, light green violin) and the closest paralog of the genes located on the hunch-specific sequence (right, dark green violin). Positive values refer to male biased expression. Genes with log_2_ fold change larger than 10 or smaller than -10 were truncated to 10 and -10 respectively.

**Fig S20.**
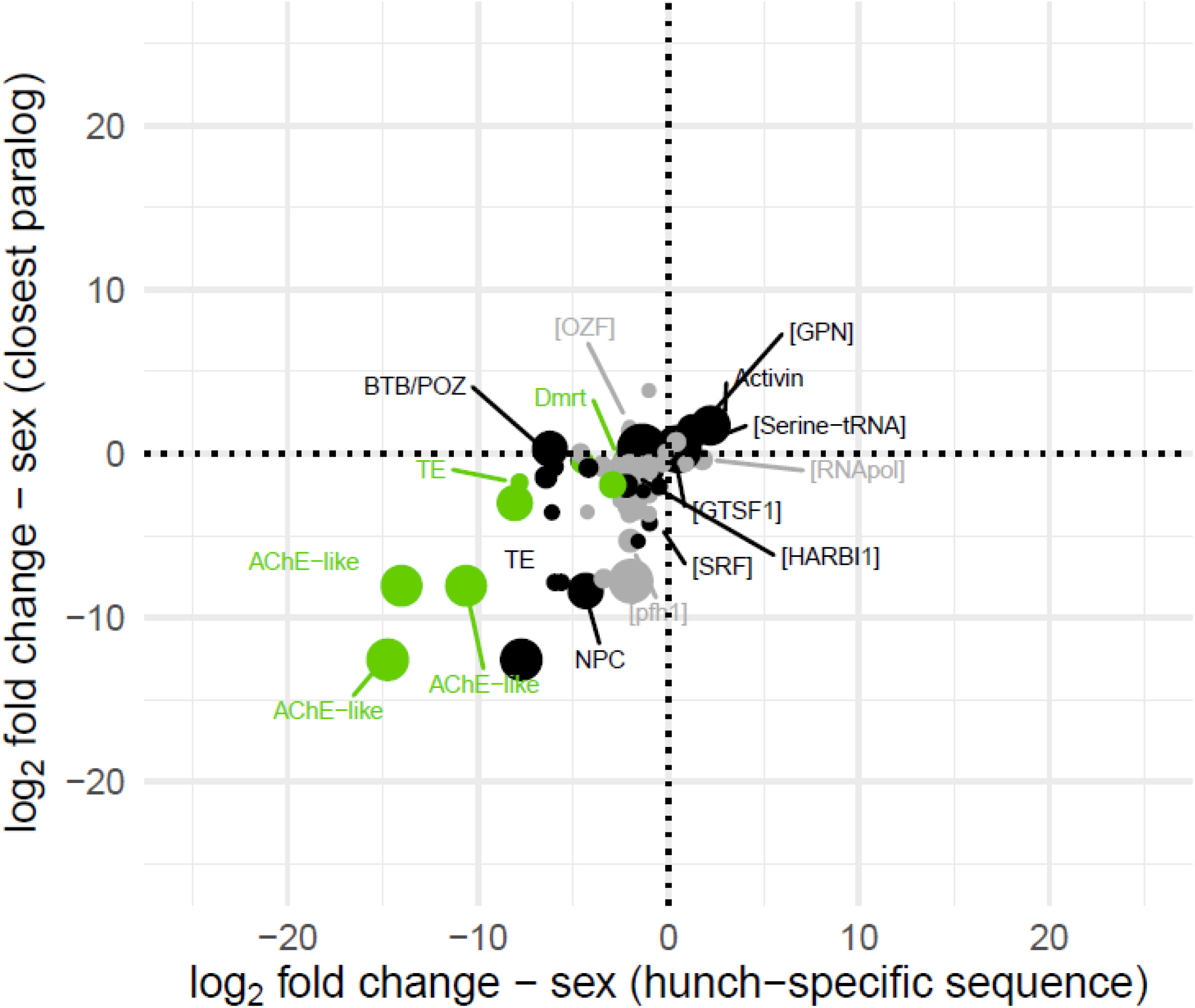
Correlation of the sex-specific expression of genes located on the hunch-specific sequence and the sex-specific expression of their closest paralog located outside the hunch-specific sequence (r_P_ = 0.53, P < 0.001). Dot size is proportional to the average normalized expression of each gene. Genes depicted in green are differentially expressed between the two male morphs and genes depicted in grey are only marginally expressed in hunched and flat males and females (normalized expression < 1).

**Fig S21.**
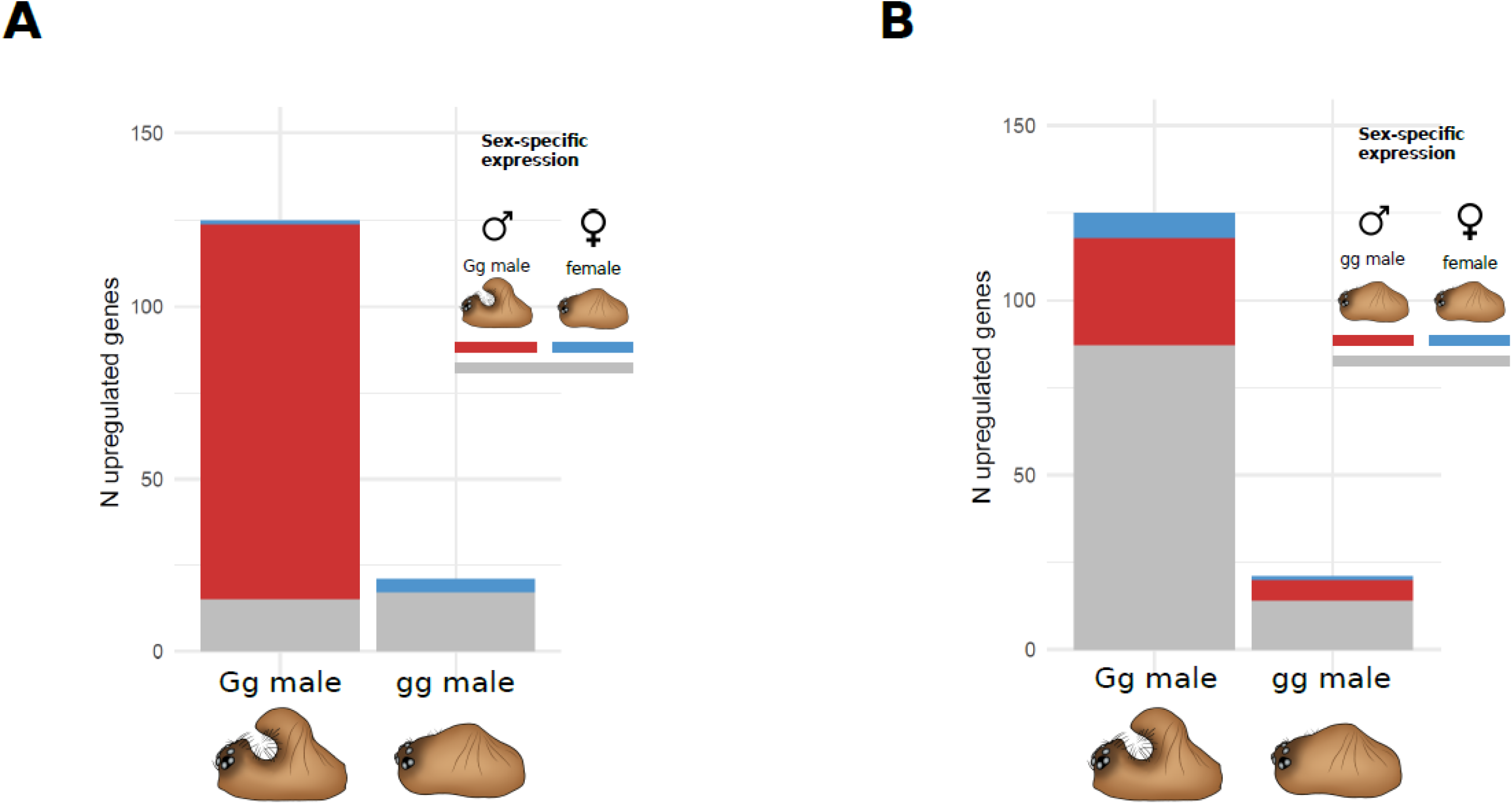
Number of differentially expressed genes between the two male morphs. Genes are color coded according to their sex-specific expression in (A) a hunched (Gg) male vs. female comparison or (B) a flat (gg) male vs. female comparison. Red genes are upregulated in males, blue genes are upregulated in females and grey genes are not differentially expressed between the sexes.

**Fig S22.**
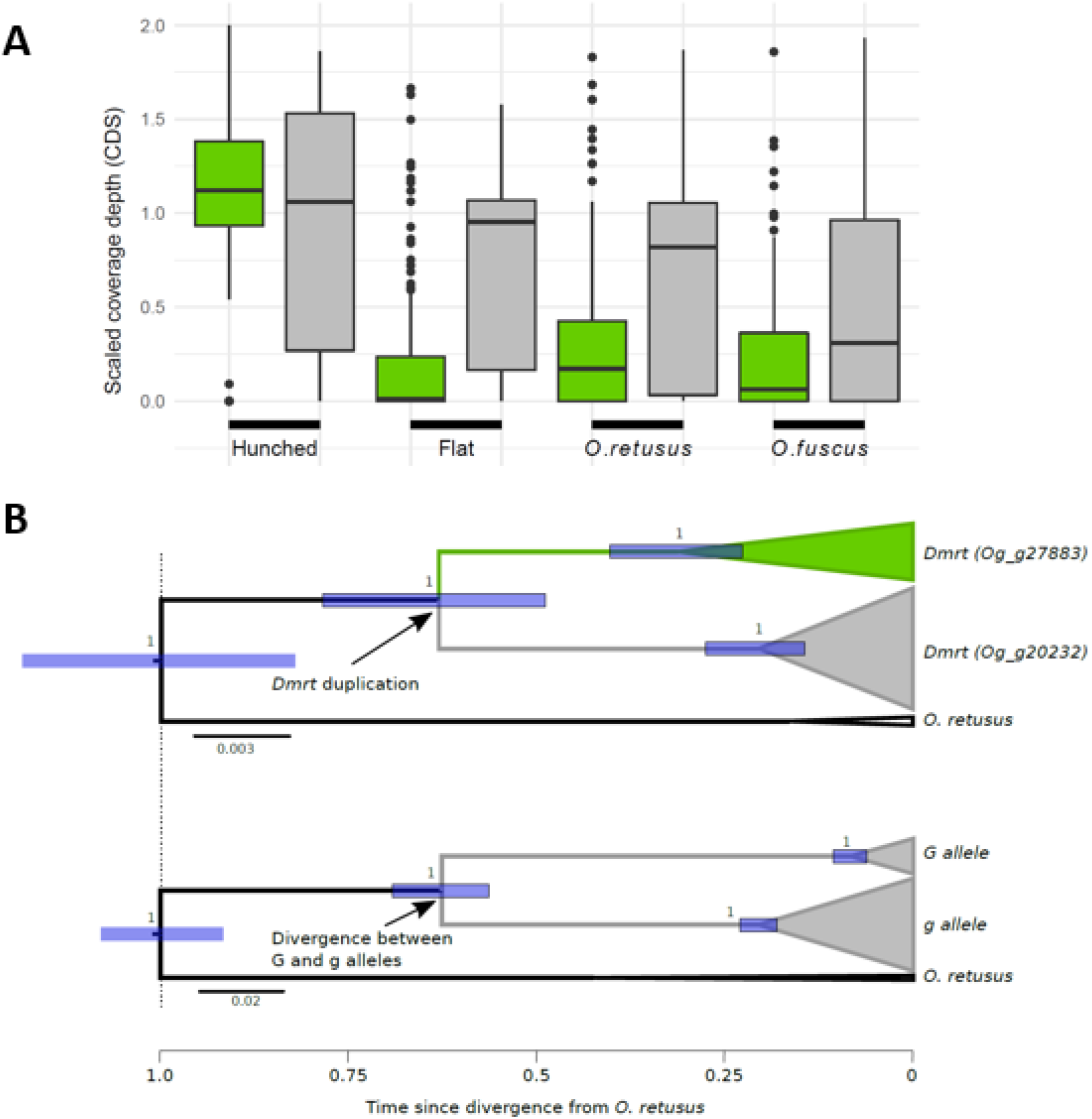
(A) Scaled coverage depths of coding sequences located on the hunch-specific sequence (green) and the closest paralogs of these coding sequences (grey) for hunched males, flat males and the closely related species *O. retusus* and *O. fuscus*. (B) Chronogram of the divergence between the *Dmrt* paralogs (above) and the divergence between the G and g allele (below). Both chronograms were scaled to the divergence from the outgroup species *O. retusus*. Triangles represent sequences within each clade. Divergence between the two alleles was based on the same genomic fragment as in Fig. 3 (ctg337: 640kb – 660kb). Node labels represent the posterior probability support of the clades. Thick blue lines represent the 95% Bayesian credible intervals of the divergence times. Midpoint rooting was used to root the trees.

